# Invasive DNA elements modify nuclear architecture by *KNOT*-Linked Silencing in plants

**DOI:** 10.1101/500496

**Authors:** Stefan Grob, Ueli Grossniklaus

**Affiliations:** Department of Plant and Microbial Biology & Zurich-Basel Plant Science Center, University of Zurich, Zollikerstrasse 107, 8008 Zurich, Switzerland

**Keywords:** 3D nuclear organization, *Arabidopsis*, gene silencing, paramutation, transgene, *KNOT*

## Abstract

**Background:** The three-dimensional (3D) organization of chromosomes is linked to epigenetic regulation and transcriptional activity. However, only few functional features of 3D chromatin architecture have been described to date. The *KNOT* is a 3D chromatin structure in *Arabidopsis*, comprising 10 interacting genomic regions termed *KNOT ENGAGED ELEMENTs* (*KEEs*). *KEEs* are enriched in transposable elements and small RNAs, suggesting a function in transposon biology.

**Results:** Here, we report the *KNOT’s* involvement in regulating invasive DNA elements. Transgenes can specifically interact with the *KNOT*, leading to perturbations of 3D nuclear organization, which correlates with the transgene’s expression: high *KNOT*-contact frequencies are associated with transgene silencing. *KNOT*-Linked Silencing (KLS) cannot readily be connected to canonical silencing mechanisms, such as RNA-directed DNA methylation and post-transcriptional gene silencing, as both cytosine methylation and small RNA abundance do not correlate with KLS. Furthermore, KLS exhibits paramutation-like behavior, as silenced transgenes can lead to the silencing of active transgenes *in trans*.

**Conclusion:** Transgene silencing can be readily connected to a specific feature of *Arabidopsis* 3D nuclear organization, namely the *KNOT*. KLS likely acts either independent or prior canonical silencing mechanisms and, hence, its characterization promises to not only contribute to our understanding of chromosome folding but moreover provides valuable insight into how genomes are defended against invasive DNA elements.

## Background

Genome organization encompasses the linear genome, the epigenome, and its 3-dimensional architecture (3D-genome). In contrast to the first two organizational levels, our understanding of the functional roles of the 3D-genome is rather poor. Chromosome conformation capture (3C) technologies [1] have facilitated its exploration, implicating it in transcriptional regulation [2], replication [3], and senescence [4]. We previously proposed a role of the 3D-genome in transposon biology in *Arabidopsis* [5]: Ten *KNOT ENGAGED ELEMENTS* (*KEEs*) (aka IHIs [6]), transposable element (TE) insertion hotspots enriched in small RNAs (sRNAs), tightly associate to form a nuclear structure termed the *KNOT* (**Fig. 1A and Additional file 1: Table S13**). The *KNOT* is conserved in plants, found in both dicots and monocots, and a potentially analogous structure may be formed by *Drosophila* piRNA clusters [5,7].

**Fig. 1.**
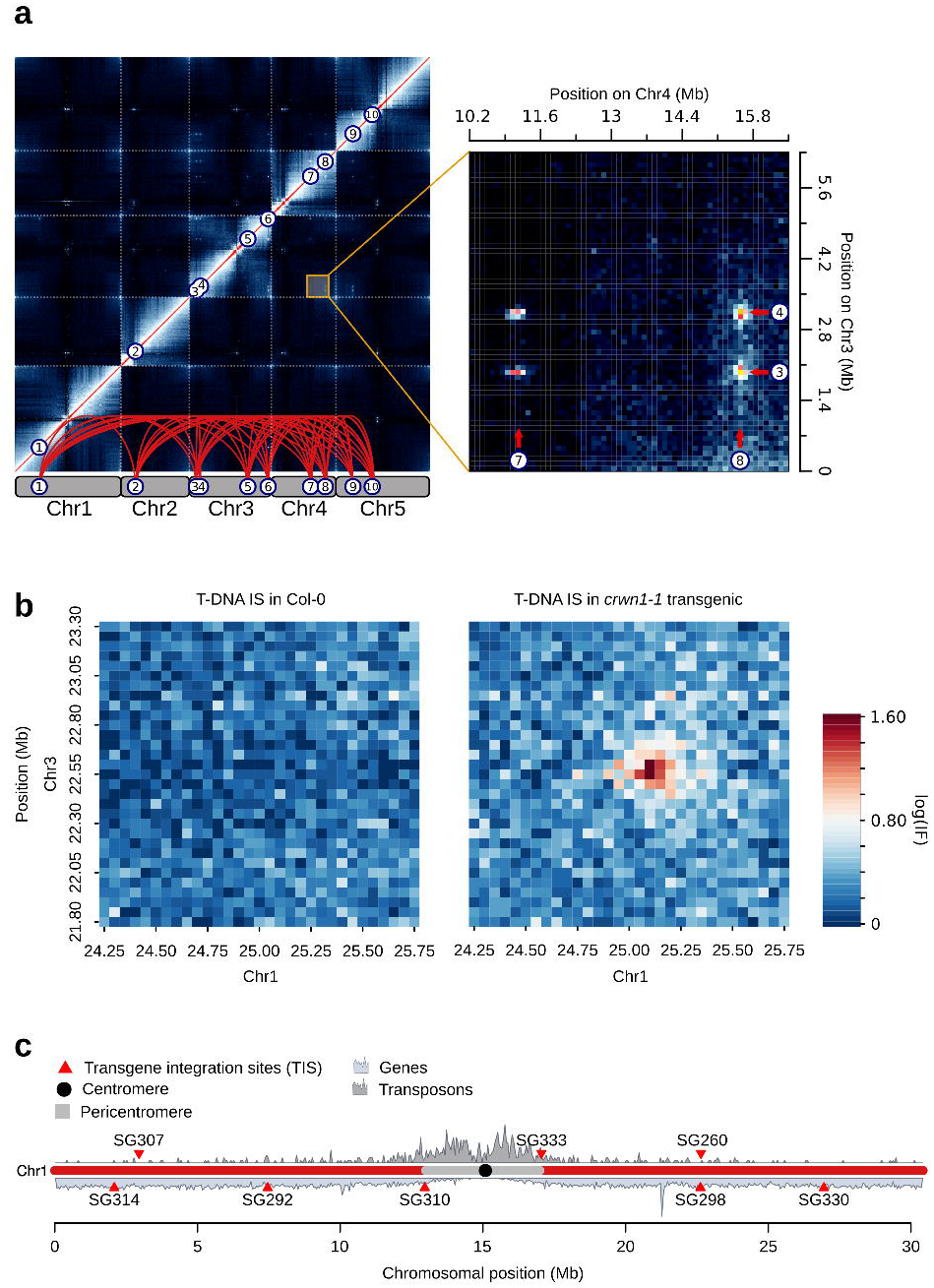
Novel *KNOT*-interactions in transgenic plants. **a)** Hi-C interactome of *Arabidopsis thaliana*. The *KNOT* is represented by network of long-range *cis*-and *trans*-contacts found between all *Arabidopsis* chromosomes (see also **Additional file 1: Table S13**). **b)** Hi-C interaction data representing interaction frequencies (IFs) between genomic regions on Chromosome 1 (Chr1) (*TIS*_*SALK* *T-DNA*_, Chr1: 25151270 – 25156323) and Chr3 (*KEE6*, 22560488 – 22580488). Ectopic IFs can be observed between the *TIS* and *KEE6*. IFs are pooled into 50 kb genomic bins (see also **Additional file 1: Figure S1A**). **c)** Representation of *TISs* on Chr1 investigated in this study. Gene and transposon density are shown to facilitate the distinction of euchromatic and heterochromatic regions.

Invasive DNA elements, such as TEs, retroviruses, and transgenes, are not only central to biotechnology but also play an important role in disease [8] and genome evolution [9]. Plants have evolved a balanced response to these elements, allowing for potential benefits, such as rapid adaptation to environmental challenges, through controlled mobility [10]. In contrast, their uncontrolled proliferation and expression, which can lead to genome instability and potentially harmful ectopic gene expression, respectively, is counteracted by the silencing of invasive elements. With transgenes, silencing has been observed since the beginning of their use (reviewed in Kooter et al., 1999) and is of concern to both, gene technology and fundamental research. In plants, many transgenic approaches are based on T-DNA vectors [12]. However, despite their common origin, vectors used to generate transgenic plants exhibit significant differences with respect to transgene expression. Certain vectors, especially those containing viral *35S* regulatory sequences [13], such as *pROK2* used to generate the insertion lines of the SALK collection [14], become more frequently silenced than others. It is unlikely that the underlying mechanism is directly associated with these transgenes, as plants must have evolved strategies to counteract invasive elements well before plant transformation was developed. Hence, although the susceptibility to silencing differs among vectors, the underlying mechanisms are likely universal irrespective of the variation with respect to silencing. The high frequency and variability of silencing among SALK lines make them an ideal system to study the control of invasive genetic elements. Suppression of such elements in plants has been associated with sRNA-mediated processes, either leading to transcript decay or DNA methylation and transcriptional silencing [13,15]. Here, we introduce an alternative silencing mechanism, *KNOT*-Linked Silencing (KLS), and show how transgenes and the 3D-genome can reciprocally influence each other.

## Results

### Ectopic 3D contacts between transgene insertion sites and the *KNOT*

We reanalyzed previously published Hi-C data [5] obtained from mutant plants and observed novel high-frequency long-range interactions that were absent in the wild type (**Fig. 1B**). In the *crwn1-1* mutant [16], caused by a T-DNA insertion, these novel interactions occur between the *CRWN1* locus and several *KEEs*. Additionally, we observed an enrichment of interaction frequencies between the transgene integration site (*TIS*) and constitutive heterochromatin of all five *Arabidopsis* chromosomes (**Fig. 1B** and **Additional file 1: Figure S1A-B**).

We hypothesized that transgene integration can induce ectopic *KEEs* that originate from the *TIS*, resulting in novel high-frequency contacts between the *TIS* and the *KNOT*. Thus, transgene integration may disturb the endogenous 3D-organization of the *TIS*. To test this, we performed 4C experiments in 8 independent, publicly available transgenic SALK lines, setting the viewpoint at the respective *TIS* (**Fig. 1C**). In parallel, we generated 4C interaction profiles of the same viewpoints in Columbia-0 (Col-0) wild-type plants, and statistically evaluated differences between transgenic and wild-type 4C profiles (**Fig. 2**). Between transgenic and wild-type lines, differential interaction analysis revealed significant differences (FDR < 0.05), predominantly coinciding with *KEEs* (6 of 8 transgenic lines) (**Fig. 2**). However, the magnitude of perturbation in the 4C profile differed considerably among lines. Three of them (SG260, SG292, and SG298) exhibited a significant change in interaction frequencies only with respect to one individual *KEE* (*KEE3* for SG292 and *KEE6* for SG260 and SG298, respectively). Other transgenic lines (SG307, SG314, and SG330) showed more severe perturbations of their 4C profile. We detected ectopic high-frequency contacts with most *KEEs* and with pericentromeric regions of all chromosomes, reminiscent of the initial observation in *crwn1-1* (**Fig. 2** and **Additional file 1: Figure S1A**). Thus, transgene integration does not solely result in the insertion of additional genetic material but can also perturb the 3D-organization of the *TIS* in a specific manner. The absence of increased *TIS*-pericentromere interactions observed in SG260, SG292, and SG298 indicates that novel *TIS-KEE* interactions are not a consequence of *TIS* dislocation towards the pericentromere.

**Fig. 2.**
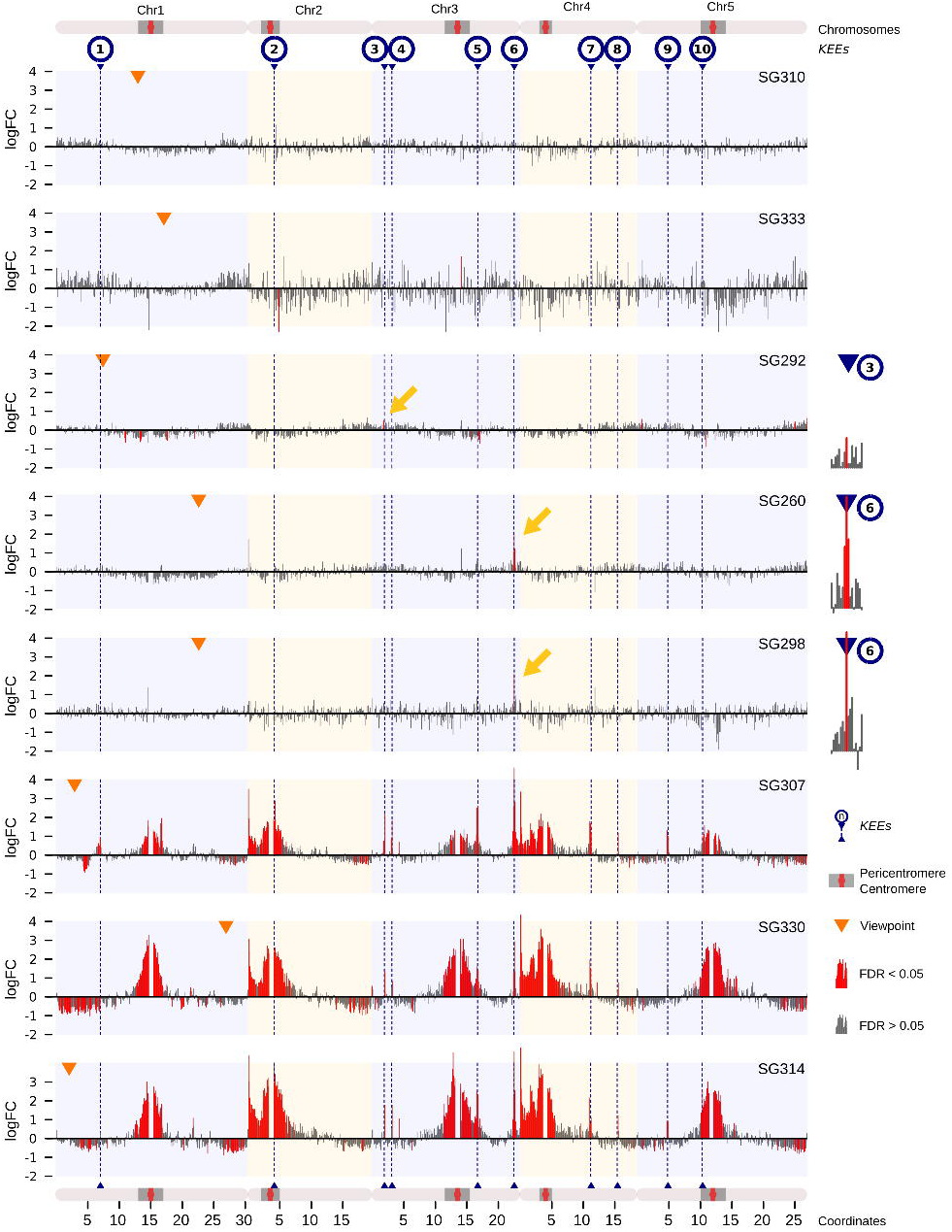
*TISs* interact with *KEEs* and pericentromeric regions. Differential analysis of 4C interactomes, including 3 wild-type and 3 transgenic 4C samples. Log2 fold changes (FC) are plotted. Grey: non-significant FC (FDR > 0.05). Red: significant FC (FDR ≤ 0.05). Orange triangles indicate viewpoints (adjacent to *TIS* on endogenous sequence). Blue triangles and dashed blue lines indicate positions of *KEEs*. Grey rectangles delineate pericentromeric regions. Interaction frequencies of single *Hind*III restriction fragments were pooled into 100 kb genomic bins. Yellow arrows indicate significant *TIS-KEE* contacts, for which magnification is given on the left.

To assess whether the ectopic *KEE6*-*TIS* interactions coincide with decreased interaction frequencies between *KEE6* and other *KEEs*, we analyzed *KEE6-KNOT* and *KEE6-CRWN1* interaction frequencies in *crwn1-1* Hi-C data and other Hi-C data sets (wild-type and transgenic) [5,17], which did not exhibit ectopic *KEE6-CRWN* interactions. Indeed, *KEE6-KNOT* interaction frequencies were decreased in *crwn1-1*, suggesting that *KEE6* is partially dislocated from the *KNOT* upon contacting the transgene (**Additional file 1: Figure S1E-F**).

### Number of insertions may influence the strength of *TIS-KNOT* interactions

To further investigate variation in the extent of 3D-genome perturbations between lines, we analyzed the number of *TIS* by Southern blotting and droplet digital PCR (ddPCR) (**Additional file 1: Table S1** and **Figure S4A**). Transgenic lines exhibiting either no significant changes in interaction frequencies, or significant alterations with respect to single *KEEs* only, harbored single insertions (SG260, SG292, SG298, SG310, and SG333). All lines that exhibited more severe alterations in 3D-organization (SG307, SG314, and SG330) carried multiple insertions. PCR-based analysis using primers flanking the insertion sites indicated that multiple copies were inserted at a single locus. However, although not observed by Southern blotting, ddPCR, and short read sequencing data (4C data), we cannot completely exclude that additional T-DNA fragments are inserted elsewhere in the genome. The occurrence of large-scale rearrangements, such as translocations, can be ruled out as we can readily detect such rearrangements by 4C (**Additional file 1: Figure S4B-D**). As half of the single-insertion and all multiple-insertion lines showed high-frequency interactions with *KEEs*, transgene copy number may influence the strength but not the potential of *TIS*-*KNOT* interactions *per se*.

### Tight *TIS-KNOT* 3D contacts coincide with transgene silencing

Next, we investigated whether ectopic *TIS*-*KEE* contacts affect the activity of the transgenes. The vector *pROK2*, used to generate the transgenic lines [14], harbors the *NPTII* kanamycin resistance gene. Thus, we visually assessed the viability of transgenic seedlings grown on medium containing kanamycin (**Fig. 3A and Additional file 1: Figure S2A**). The phenotypes were uniform in three distinct populations per transgenic genotype, stressing the robustness of the transcriptional state of the transgenes (**Additional file 1: Figure S2A**). Viability significantly anti-correlated with *TIS*-*KEE* interactions and was strongly reduced in lines with the highest *KNOT* interactions, phenocopying the absence of *NPTII* in the wild type (**Fig. 3A**-**B**). Lines with significantly increased interaction frequencies with the *KNOT* but not the pericentromeres as well as lines without increased *KNOT* interaction frequencies were not significantly affected by kanamycin, thus showing sufficient *NPTII* expression (**Fig. 3A**). We confirmed these results by RNA sequencing data, which revealed a significant anti-correlation between *TIS*-*KEE* interaction frequencies and *NPTII* expression (**Fig. 3C**) that itself significantly correlated with viability on kanamycin (**Fig. 3D**). As the strength of *TIS*-*KNOT* interactions negatively correlated with *NPTII* expression, we propose an involvement of the *KNOT* in transgene silencing.

**Fig. 3.**
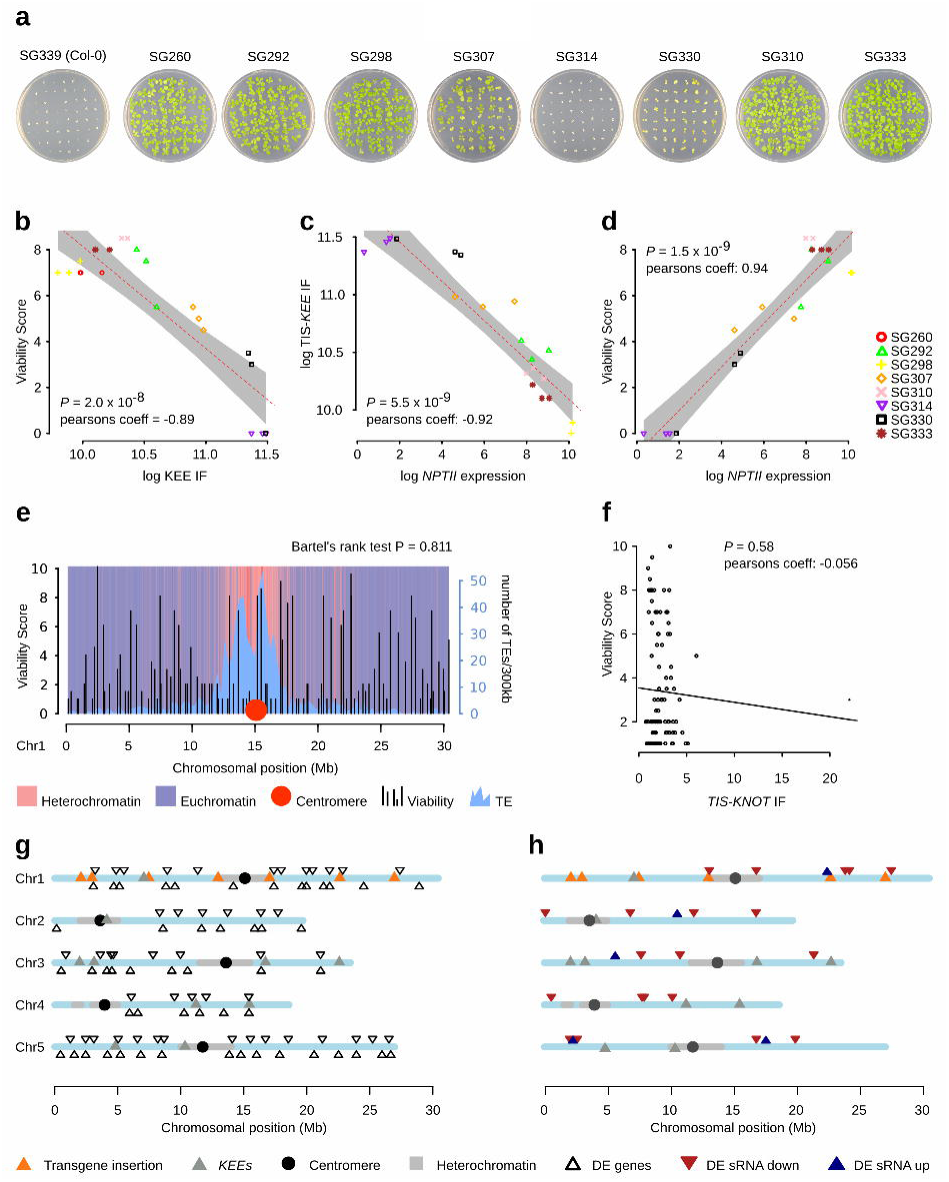
Transgene silencing by *KNOT*-mediated silencing. **a)** Seedlings growing on medium containing kanamycin show variable resistance (see also Additional file 1: **Figure S2A)**. **b)** Pearson’s correlation between phenotypically assessed viability in presence of kanamycin and *TIS* IFs with *KEEs* and pericentromeres. **c)** Pearson’s correlation between *NPTII* transgene expression and *TIS* IFs with *KEEs* and pericentromeres. **d)** Pearson’s correlation between *NPTII* transgene expression and phenotypically assessed kanamycin resistance. **e)** Viability score (10 - fully viable, 0 - dead) of transgenic seedling populations (n = 30) grown on selective medium. Transgenic lines were selected by randomly choosing a homozygous SALK line (www.signal.salk.ed/cgi-bin/homozygots.cgi) for each 300 kb genomic bin on Chr1. Numbers of transposons are indicated as a proxy for the presence of heterochromatin. Euchromatic and heterochromatic regions (purple and light red) correspond to chromatin states 1-7 and chromatin states 8-9, respectively, as previously defined [34]. **f)** Pearson’s correlation analysis of IFs between the prospective *TIS* and the *KNOT* in the wild type and the viability score of transgenic lines with insertions at the respective TIS. *TIS-KNOT* IFs were calculated from Col-0 wild-type Hi-C matrices (100 kb bins) [5]. **g)** Differentially expressed genes between Col-0 wild-type and combined expression data of all transgenic lines. For each line RNA sequencing was performed in triplicate (see also **Additional file 1: Figure S2B**). **h)** Differential analysis of sRNA-seq data. Genomic bins (500 bp) exhibiting significant changes (logFC > 2; FDR < 0.01) between S-and A-lines (see also **Additional file 1: Figure S3A**).

Interestingly, repressive genomic neighborhoods of the *TISs* did not appear to affect either transgene expression or associated perturbations in *TIS* 3D-organization: transgenes inserted into constitutive heterochromatin (SG310 and SG333) (**Fig. 1C**) were neither silenced nor exhibited strong 3D-perturbations, whereas certain *TISs* in euchromatin showed significant perturbations and were silenced. To corroborate this observation, we grew 99 homozygous SALK lines carrying insertions distributed along chromosome 1 on selective medium and scored their viability associated with *NPTII* expression. We did not observe decreased viability of lines that carry transgenes in repressive heterochromatin (**Fig. 3E**). Moreover, statistical analysis rejected a non-random distribution of viability scores, a finding supported by a previous study [18]. We cannot exclude that upon transformation, chromosomal localization may have influenced transgene expression, leading to counterselection of T-DNAs inserted into repressive environments. However, as they would not have been retrieved otherwise, all transgenes analyzed here were initially expressed and acquired a distinct expression state since. Hence, our results suggest that at least *de novo* silencing of transgenes is independent of the epigenetic environment of the *TIS*.

Furthermore, transgene silencing cannot be predicted based on wild-type interaction frequencies of a prospective *TIS* and the *KNOT*. Using Hi-C data from wild-type plants [5], we did not observe a significant correlation between the interaction frequencies of the prospective *TIS* with the *KNOT* and transgene silencing (**Fig. 3F**), indicating that the 3D-organization of the prospective *TIS* does not predispose for silencing.

To investigate whether perturbing the 3D-organization of the *TIS* is limited to transgene expression or whether *TIS*-*KNOT* contacts also affect neighboring endogenous gene expression, we performed RNA sequencing. We analyzed triplicate mRNA from seven lines to test whether expression of genes surrounding the *TIS* differed between wild-type and transgenic lines, indicative of an effect of novel *TIS*-*KNOT* interactions. We found that transcriptional silencing is restricted to the transgene, as there was no enrichment of differentially expressed genes in the neighborhood of the *TIS* or the *KEEs* (**Fig. 3G** and **Additional file 1: Figure S2B**).

Endogenous loci evade KLS, indicating specificity to invasive genetic elements. Furthermore, although the genomic region encompassing the *TIS* and nearby genes is folded into a repressive environment, silencing is limited to the transgene itself. Thus, a perturbation of nuclear architecture alone is not sufficient to silence gene expression and other, yet to be discovered, factors may play a role in KLS specificity.

### Transgene silencing does not require DNA methylation

Next, we aimed at putting KLS into the context of established silencing mechanisms in plants. There have been numerous previous reports on transgene silencing and the underlying mechanisms have been deciphered [13]. Two principle mechanisms are proposed to initiate and/or maintain transgene silencing: transcriptional gene silencing (TGS) and post-transcriptional gene silencing (PTGS) lead to transcriptional arrest and mRNA degradation, respectively [19]. Homology-dependent gene silencing, another term often used for transgene silencing, can depend on either TGS [20] or PTGS [21]. It can lead to simultaneous silencing of various homologous sequences and, hence, exhibits *trans*-silencing effects [22,23]. SALK T-DNA lines were found to be subjected to TGS, involving the accumulation of promoter-specific sRNAs and elevated levels of cytosine methylation (mC) in transgene promoters, mediated by the RNA-directed DNA-methylation (RdDM) pathway [23].

We first investigated mC levels in the nopaline synthase promoter (*nosP*) driving *NPTII* in three active (A-lines) and three silenced lines (S-lines) by Sanger sequencing after bisulfite conversion (**Fig. 4C**). In average, S-lines showed elevated mC levels and weak correlations with both, kanamycin sensitivity and *KEE* interaction frequencies (**Fig. 4C-E** and **Additional file 1: Figure S2C-F**). However, SG314, exhibiting significantly higher mC levels than all other lines, had a major effect on the statistical analysis. By omitting SG314, no significant mC enrichment in S-lines and no significant correlation between either transgene silencing or *KEE* interaction frequencies and mC levels was observed (**Fig. 4C-E**). In all transgenic lines, including SG314, the overall *nosP* mC levels were lower than expected for RdDM and comparable or below average genomic mC levels [24–27]. In summary, although one transgenic line (SG314) exhibited elevated mC levels, which may be associated with RdDM, other silenced lines (SG307 and SG330) showed low mC levels, indistinguishable from active transgenes. Therefore, mC is not necessary for the silencing of the investigated transgenes and we conclude that mC-dependent TGS, such as RdDM, is not a prerequisite for KLS. Consistent with these results, mC-independent transcriptional gene silencing has previously been reported [28].

**Fig. 4.**
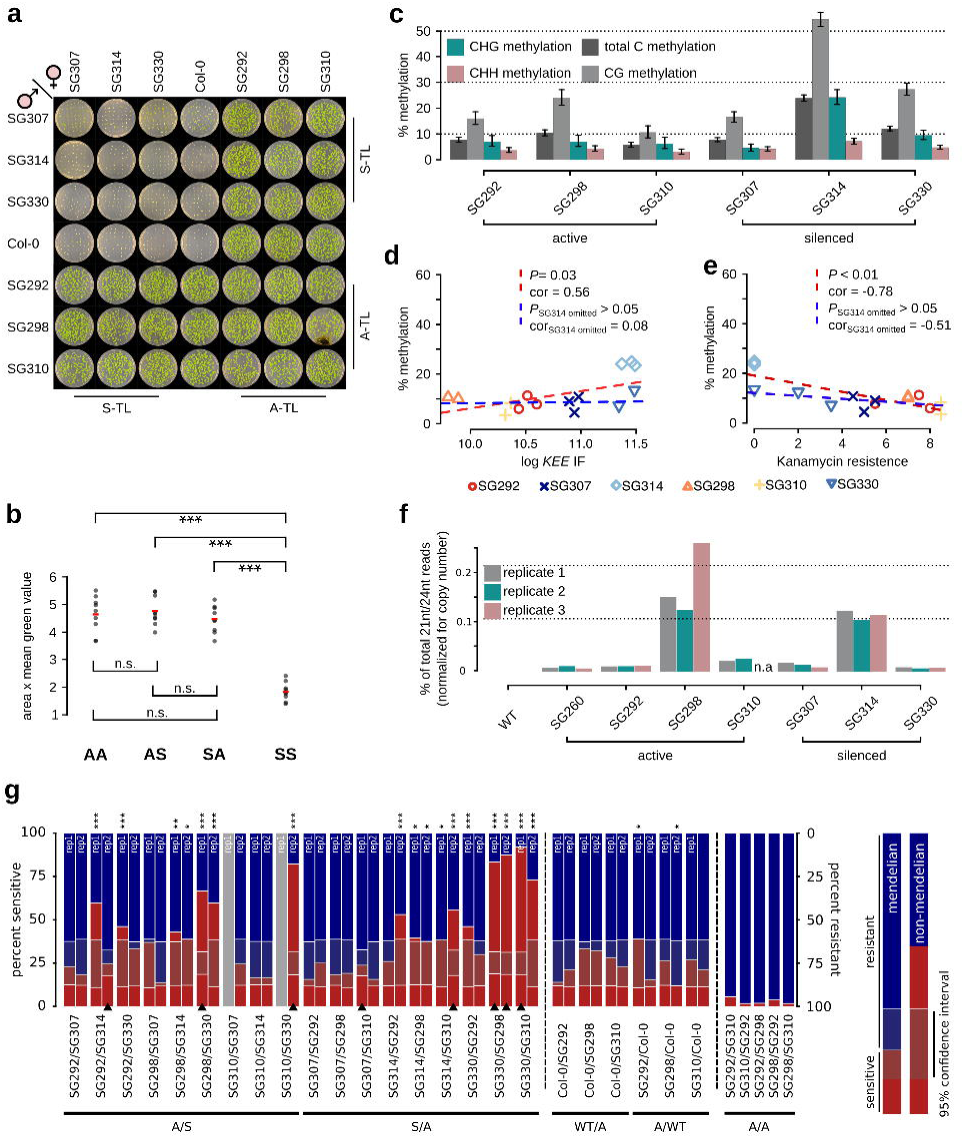
KLS is independent of canonical silencing pathways. **a)** Reciprocal crosses between silenced and active transgenic lines. Images were acquired from 14-day-old seedlings. **b)** Area and mean “green” value were assessed by ImageJ. Student’s *t*-tests were performed to assess significant differences between all quarters (SS, AA, AS, SA) of the diallel cross (*FDR*_*SSvsAA*_ = 2.7×10^−7^, *FDR*_*SSvsSA*_ = 1×10^−8^, *FDR*_*SSvsAS*_ = 1×10^−8^, *FDR*_*AAvsSA*_ = 0.87, *FDR*_*AAvsAS*_ = 0.88, *FDR*_*SAvsAS*_ = 0.47) (**Additional file 1: Table S6)**. **c)** Bisulfite Sanger sequencing of *nosP* (301 bp on 3’-end). Methylation levels in all contexts significantly differed between active and silenced lines and also between individual lines (**Additional file 1: Table S4, Figure S2C-F**). Error bars: Wilson 95% confidence intervals. **d)** Pearson’s correlation analysis between 4C IFs with *KEEs* and pericentromeres (*KEE*-IF) and *nosP* methylation levels. Weak correlation was observed (red line). Non-significant correlation was observed when the highest methylated line (SG314) is omitted (blue line). **e)** Correlation between kanamycin resistance phenotype and *nosP* methylation levels. Weak correlation was observed (red line). Non-significant correlation was observed when SG314 was omitted (blue line). **f)** Percentage of 21nt and 24nt sRNA-seq reads found within *pROK2*. For each genotype, biological triplicates were assessed (number of reads were normalized by transgene copy number) (**Additional file 1: Table S1**). **g)** Segregation in F2 seedlings. Chi-square tests were performed to test for deviation from Mendelian segregation (Null-hypothesis: 0.25/0.75 (sensitive/resistant), * 0.05 > *P* ≥ 0.01, ** 0.01 > *P* ≥ 0.001, *** *P* < 0.001). Confidence interval indicates the range, in which Mendelian segregation cannot be rejected. Bars with black triangles stem from pooled data of 4 individual F1 siblings (n = up to 4 x 52 seedlings), non-marked bars stem from mixed seeds of the 4 siblings (n = up to 52 seedlings). Grey bars: data not available (**Additional file 1: Table S7, Table S8, Figure S3C**).

### sRNA abundance does not correlate with KLS

To assess a possible involvement of sRNAs in KLS, we conducted sRNA sequencing (sRNA-seq). First, we analyzed the abundance of sRNAs mapping to the *pROK2* transgene. In case of a significant involvement of sRNAs in silencing the investigated transgenes and, thus, KLS, we expected to find high levels of associated sRNAs in S-lines and low levels in A-lines. We detected sRNAs associated with *pROK2* in all transgenic lines, although to variable extents (**Fig. 4F** and **Additional file 1: Figure S3B**). In accordance with our DNA methylation analysis, sRNAs were abundant in SG314, yet, no general correlation between sRNA levels and transgene silencing was found. By normalization of sRNA reads to transgene copy number, an A-line (SG298) even exhibits the highest abundance of sRNAs in all analyzed lines (**Additional file 1: Figure S3B**). Additionally, both the silenced lines SG330 and SG307 showed indistinguishable sRNA levels from A-lines (SG260, SG292, SG310). We conclude that sRNAs are neither sufficient nor necessary to silence these transgenes. In summary, our findings suggest that neither DNA methylation nor sRNAs play a primary role in silencing the investigated transgenes and that KLS does not depend on RdDM-related TGS.

To perform a genome-wide analysis of sRNA abundance in the investigated lines, sRNA reads were binned to 500 bp genomic regions and subsequently analyzed to detect loci of differential sRNA association (**Additional file 1: Figure S3A**). The sRNA profiles of active and silenced transgenic lines were very similar, identifying only few distinct differential loci (**Fig. 3H**). An analysis of genomic features overlapping the identified differential loci did not reveal obvious candidate factors involved in transgene silencing. We subsequently compared the identified differential sRNA loci with differentially expressed genes obtained from the mRNA-seq experiment using the same contrast (active vs. silenced transgenic lines) and no overlap between the two data sets was found. Similarly, analysis of the differentially expressed genes of this contrast did not provide candidates associated with transgene silencing. Our results suggest that sRNAs do not appear to be directly involved in silencing the investigated transgenes.

By performing an alternative experiment, we aimed to independently confirm that sRNAs are not a prerequisite of KLS. Specifically, we used a genetic approach to test whether PTGS is involved in KLS. As PTGS involves sRNAs that lead to mRNA decay, it can silence genes in *trans*. Thus, the progeny of a cross between an S-and an A-line should be at least partially silenced, as transgenes identical in sequence are present in both parental lines, such that mRNA from both transgenes should be affected by the same sRNAs. We performed reciprocal crosses using seven parental lines: one wild-type, three S-(SG307, SG314, SG330), and three A-lines (SG292, SG298, SG310) (**Fig. 4A**). This resulted in 8 progeny groups, either derived from two S-lines (SS), two A-lines (AA), two groups of progenies with parents of converse transcriptional state (SA and AS), and 4 groups of hemizygous transgenic progeny. We assessed their viability reflecting *NPTII* expression by growing F1 seedlings on selective medium and measuring the area and mean green fraction intensity of imaging data (**Fig. 4A-B**). The transgene expression state behaved as a heritable dominant trait (**Fig. 4A**). SS progeny, lacking *NPTII* expression, exhibited significantly reduced viability compared to all other groups, whereas as SA, AS, and AA groups did not significantly differ from each other (**Fig. 4B**). Thus, in F1 seedlings, KLS behaves as a recessive trait with Mendelian inheritance. This excludes the involvement of diffusible sRNAs acting in *trans*, suggesting that PTGS is unlikely involved in KLS.

### KLS shows paramutation-like features

To assess whether the F2 generation also follows Mendelian segregation, we cultivated progeny of the above-described crosses on non-selective medium and allowed four plants of each F1 population to self-fertilize. We then analyzed the segregation in response to kanamycin in the F2 seedling populations. Assuming Mendelian segregation, double hemizygous F1 plants containing a silenced and active *NPTII* transgene are expected to produce 25% kanamycin-sensitive offspring (**Additional file 1: Figure S3E**). Employing PCR-based genotyping, we could confirm genetic Mendelian segregation for both transgenes (**Additional file 1: Table S9**). However, phenotypically we observed a deviation from Mendelian segregation in a large fraction of the F2 populations, manifested in significantly higher proportions (up to 92%) of kanamycin-sensitive seedlings (**Fig. 4G and Additional file 1: Figure S3C**). The observed phenotypic segregation distortion indicates that a large fraction of parentally active transgenes underwent *de novo* silencing, a process reminiscent of paramutation [29]. In support of our observation, *trans*-silencing between transgenes has been observed before [22,23]. Importantly, during the entire crossing procedure, the *trans*-silencing effect depends on the initial presence of a silenced transgene, as F2 seedling populations derived from AA crosses did not exhibit *trans*-silencing phenotypes (**Fig. 4G**). However, genotyping and subsequent quantification of *NPTII* transcripts by ddPCR of single F2 plants revealed that the presence of the paramutagenic allele is not necessary for the paramutagenic effect in the F2 generation, as plants derived from AS crosses, which were homozygous for the A but lacking the S transgene, still exhibited full *NPTII* silencing (**Additional file 1: Figure S3D**). The observed proportions of silencing also exclude a potential dosage effect of diffusible sRNAs associated with post-transcriptional gene silencing that are produced by the parentally silenced transgene (**Additional file 1: Figure S3E-F**). In summary, transgenes silenced by KLS show a paramutation-like behavior, but their initial silencing is not correlated with mC and sRNAs targeting the transgene, indicating a novel mechanism depending on 3D-genome interactions.

## Discussion

### The *KNOT* is a novel player of the genome’s defense system

Our results suggest that insertion of transgenes has more profound effects on genome structure than previously anticipated, as not only genetic material is added, but also the 3D-architecture of the *TIS* can be severely perturbed. These alterations have a profound impact on the transgenes themselves, as architectural perturbations can clearly be associated with the expression state of the transgenes. Importantly, the observed perturbations are not random. Moreover, we detected specific ectopic interactions with the *KNOT*, suggesting its involvement in the nuclear defense system against invasive genetic elements.

### KLS does not depend on canonical silencing pathways

In our studies on the nature of KLS, we could not find strong evidence for an involvement of either PTGS or canonical TGS, suggesting that KLS is at least initially independent of these silencing mechanisms. In support, a previous study showed that the number of *KEE*s is not reduced in mutants leading to the de-repression of silenced genes [6]. The epigenetic marks affected in these mutants include repressive histone modifications, such as H3K27me3 (*clf;swn* double mutant) and H3K9me2 (*suvh4;suvh5;suvh6* triple mutant), DNA methylation (*ddm1, met1, cmt3*), and epigenetic processes affecting silencing by other means (*mom1*). This suggests that epigenetic marks commonly associated with gene silencing, such as H3K9me2, H3K27me3, and mC, are not necessary for interactions among *KEEs*. Hence, an involvement of these canonical repressive marks in the recruitment of T-DNAs to the *KNOT*, thereby initiating KLS, is unlikely. Interestingly, in many of these mutants an identical set of ectopic *KEEs* can be observed in apparently pre-defined positions, which show a significant enrichment of *VANDAL6* and *ATLANTYS3* TEs, both of which are highly enriched in the ten canonical *KEEs* (**Additional file 1: Figure S3G**). This finding suggests that inactive *KEE* regions exist in the genome, whose functional activation may rely on active transcription of TEs.

### KLS is a dynamic process

We observed that *TIS-KNOT* interactions alone are insufficient for transgene silencing, which only occurs in lines that also acquired high-frequency *TIS*-pericentromere interactions. We hypothesize that *TIS-KNOT* interactions may initiate transgene silencing, which could then be followed by a secondary alteration of the *TIS*’ 3D-organization, leading to a tight association with constitutive heterochromatin and complete silencing of the transgene. Although not yet observed at the time of writing, continuous growing of lines showing exclusively *TIS-KNOT* interactions, such as SG298 and SG260, over several generations may corroborate this hypothesis.

KLS shows a paramutation-like behavior, whereby the transcriptional state of one transgene can be transferred to another. The KLS *trans*-silencing activity differs from classical paramutation, as it affects non-homologous loci and seems to depend on passage through an additional generation, the latter having also been observed for other transgenes with paramutation-like behavior [30]. Furthermore, the maintenance of the repressed paramutated state in maize requires factors involved in the biogenesis of 24 nt long sRNAs, which are homologous to components of the RdDM pathway in *Arabidopsis* [31]. Similarly, sRNAs have previously been implicated in homology-dependent *trans*-silencing in *Arabidopsis* [22,23]. In contrast to these findings, sRNAs do not seem to play a determining role in KLS, making their involvement in KLS-related *trans*-silencing unlikely.

Transgene silencing represents an acquired epigenetic state, which is stably inherited over subsequent generations. All the transgenic lines analyzed here initially exhibited active *NPTII* transcription [14]; hence, KLS is a dynamic process, potentially established and augmented over consecutive generations. Previous reports on transgenerational epigenetic inheritance implicated DNA methylation in this process [32,33]. Our results suggest an independent role for 3D-genome organization in the transgenerational epigenetic inheritance of silenced transgenes. Although we observed tight *TIS-KEE* interactions stably over subsequent generations, we also show that KLS may contribute to the plasticity of transgenerational epigenetic inheritance through a paramutation-like *trans*-silencing mechanism.

### Molecular mechanism of KLS remains to be deciphered

Very likely, KLS involves a set of protein cofactors that mediate 3D *TIS-KEE* interactions. The identification of these cofactors will be essential for a better understanding of KLS and its embedding within other nuclear processes. However, this search will be challenging due to the technical inaccessibility of KLS phenotypes, such as *TIS-KEE* interactions, for large-scale genetic screening.

KLS represents a previously uncharacterized mechanism to defend the genome against invasive DNA elements. Hence, KLS is not only important for a basic understanding of gene regulation in the context of the 3D-genome but is also of great interest to plant biotechnology, as transgene integration may have a larger impact on genome architecture than previously thought.

## Conclusions

Mobile invasive DNA elements can threaten proper genome function. Hence, their transcription is regulated and can be shut down by cellular processes known as gene silencing mechanisms. We here present a novel aspect of gene silencing, which is linked to the *KNOT*, a specific 3D-chromosomal structure. Our results suggest a functional role of 3D-genome folding in the defense against invasive elements. KLS appears to be independent of previously published silencing mechanism, whose hallmarks are increased DNA methylation and RNA interference. In fact, KLS may even underlie these silencing mechanisms. Interestingly, the *KNOT* is conserved within the plant kingdom; thus, KLS may represent a basal silencing mechanism common to most plants.

## Methods

Detailed description on experimental procedures, materials used, and statistical analysis are available in **Supplemental Materials**.

## List of abbreviations

*KEE*: *KNOT* engaged elements
3D: three-dimensional
sRNA: small RNA
KLS: *KNOT*-linked silencing
TE: transposable element
TGS: transcriptional gene silencing
PTGS: post-transcriptional gene silencing
RdDM: RNA-dependent DNA methylation
IF: interaction frequency
TIS: transgene insertion site
mC: methyl cytosine
A-line: active transgenic line
S-line: silenced transgenic line
sRNA-seq: sRNA sequencing
*nosP*: nopaline synthase promotor
ddPCR: droplet digital PCR
4C: circular chromosome conformation capture

## Declarations

### Availability of data and material

4C and RNA, and sRNA sequencing data are publicly available at the Short Read Archive (SRA; https://www.ncbi.nlm.nih.gov/sra/) under accession SRP126992. Codes for data processing and analysis are available upon request.

### Competing interests

The authors declare no competing financial interests.

### Funding

This work was supported by the University of Zurich and an Advanced Grant of the European Research Council (MEDEA-250358) to U.G.

### Authors’ contributions

S.G and U.G. conceived the study; S.G. designed and performed the experiments, analyzed the data, and wrote the manuscript; U.G. acquired funding and helped with data interpretation and writing of the manuscript.

## Acknowledgements

We thank F. Fiscalini for help with genotyping, V. Gagliardini for setting up ddPCR assays, A. Hermann for help with crosses, A. Sarazin for advice on sRNA-seq, C. Eichenberger, A. Bolaños, D. Guthörl, A. Frey, and P. Kopf for general laboratory support, P. Jullien for viability scoring, and P. Jullien, J. Vermeer, Ö. Kartal, H. Vogler, and M. Schmid for valuable input on the manuscript.

