## Supplemental Materials for "Invasive DNA elements modify nuclear architecture by *KNOT*-Linked Silencing in plants"

### Supplementary Tables

**Table S1.** Transgene copy number analyzed by droplet digital PCR (ddPCR). The transgene copy number was assessed using the *NPTII* (*KAN*) gene relative to two endogenous single copy loci (*FIE* (AT3G20740) and *LYS* (AT5G62150)). The final copy number was determined by the mean of *KAN/FIE* and *KAN/LYS* ratios.

| **Parental** | **F1** | ***KAN* rep1** (copies/µl) | ***KAN* rep2** (copies/µl) | ***FIE* rep1** (copies/µl) | ***FIE* rep2** (copies/µl) | ***LYS* rep1** (copies/µl) | ***LYS* rep2** (copies/µl) | ***KAN/FIE*** | ***KAN/LYS*** | **Copy number** |
|---|---|---|---|---|---|---|---|---|---|---|
| SG261 | SG261 | 55.9 | 62.9 | 59.1 | 53.8 | 61.1 | 54.3 | 1.1 | 1 | 1 |
| SG261 | SG368 | 36.2 | 35.1 | 38 | 39.3 | 39.8 | 36.4 | 0.9 | 0.9 | 1 |
| SG261 | SG371 | 87 | 82.8 | 96.6 | 107.7 | 91.8 | 108.7 | 0.8 | 0.8 | 1 |
| SG292 | SG335 | 52.3 | 49.5 | 50.5 | 50.3 | 54 | 49.1 | 1 | 1 | 1 |
| SG292 | SG337 | 132 | 144.7 | 145 | 148 | 145.6 | 159 | 0.9 | 0.9 | 1 |
| SG292 | SG369 | 170 | 146 | 169 | 168 | 177 | 182 | 0.9 | 0.9 | 1 |
| SG298 | SG350 | 298 | 302 | 324 | 319 | 337 | 309 | 0.9 | 0.9 | 1 |
| SG298 | SG361 | 168 | 161 | 160 | 171 | 184 | 175 | 1 | 0.9 | 1 |
| SG298 | SG362 | 61.9 | 61.7 | 51.1 | 42.8 | 43.7 | 47.6 | 1.3 | 1.4 | 1 |
| SG307 | SG342 | 304 | 284 | 90.1 | 84 | 82.9 | 78.9 | 3.4 | 3.6 | 4 |
| SG307 | SG355 | 346 | 370 | 139 | 119 | 85.6 | 128 | 2.8 | 3.4 | 3 |
| SG307 | SG356 | 408 | 494 | 144.7 | 176 | 159 | 206 | 2.8 | 2.5 | 3 |
| SG310 | SG340 | 127.7 | 115 | 134 | 133 | 135 | 126 | 0.9 | 0.9 | 1 |
| SG310 | SG358 | 37.9 | 40.7 | 38.7 | 41.3 | 40.4 | 42.5 | 1 | 0.9 | 1 |
| SG314 | SG346 | 628 | 576 | 215 | 215 | 217 | 258 | 2.8 | 2.5 | 3 |
| SG314 | SG354 | 562 | 499 | 166 | 187 | 183 | 196 | 3 | 2.8 | 3 |
| SG314 | SG357 | 355 | 361 | 116.9 | 125.8 | 133.3 | 129.3 | 3 | 2.7 | 3 |
| SG330 | SG352 | 472 | 484 | 94 | 103.2 | 107.8 | 103.4 | 4.8 | 4.5 | 5 |
| SG330 | SG353 | 926 | 843 | 134.8 | 139 | 139 | 153 | 6.5 | 6.1 | 6 |
| SG330 | SG359 | 1334 | 1390 | 275 | 246 | 273 | 281 | 5.2 | 4.9 | 5 |
| SG333 | SG366 | 70.8 | 60.9 | 61.9 | 69.5 | 76.8 | 76.2 | 1 | 0.9 | 1 |
| SG333 | SG367 | 50.2 | 51.6 | 55.2 | 53.4 | 63.8 | 63.5 | 0.9 | 0.8 | 1 |
| SG333 | SG373 | 35.4 | 36.7 | 43.8 | 41 | 42.8 | 44 | 0.9 | 0.8 | 1 |
| SG339 | SG339B | 0.2 | 0.07 | 164 | 168 | 171 | 60.2 | 0 | 0 | 0 |
| SG339 | SG339C | 0.07 | 0.15 | 105.9 | 105 | 112 | 111.8 | 0 | 0 | 0 |

**Table S2.** Kanamycin resistance score. Viability phenotype (scale 0 (dead) to 10 (no effect of Kanamycin)) was scored double blindly based on images acquired from 14-day-old seedlings grown on kanamycin containing medium.

| Parental | F1 | score 1 | score 2 | mean score |
|---|---|---|---|---|
| SG261 | SG261 | 7 | 7 | 7 |
| SG261 | SG368 | 7 | 7 | 7 |
| SG261 | SG371 | 7 | 7 | 7 |
| SG292 | SG335 | 5 | 6 | 5.5 |
| SG292 | SG337 | 8 | 8 | 8 |
| SG292 | SG369 | 7 | 8 | 7.5 |
| SG298 | SG350 | 7 | 7 | 7 |
| SG298 | SG361 | 8 | 7 | 7.5 |
| SG298 | SG362 | 7 | 7 | 7 |
| SG307 | SG342 | 5 | 6 | 5.5 |
| SG307 | SG355 | 5 | 5 | 5 |
| SG307 | SG356 | 4 | 5 | 4.5 |
| SG310 | SG340 | 9 | 8 | 8.5 |
| SG310 | SG358 | 9 | 8 | 8.5 |
| SG314 | SG346 | 0 | 0 | 0 |
| SG314 | SG354 | 0 | 0 | 0 |
| SG314 | SG357 | 0 | 0 | 0 |
| SG330 | SG352 | 3 | 3 | 3 |
| SG330 | SG353 | 3 | 4 | 3.5 |
| SG330 | SG359 | 0 | 0 | 0 |
| SG333 | SG366 | 8 | 8 | 8 |
| SG333 | SG367 | 8 | 8 | 8 |
| SG333 | SG373 | 9 | 7 | 8 |
| SG339 | SG339A | 0 | 0 | 0 |
| SG339 | SG339B | 0 | 0 | 0 |
| SG339 | SG339C | 0 | 0 | 0 |

**Table S3.** Methylation analysis by Sanger sequencing after bisulfite treatment. Target: Chloroplast DNA. Chloroplast DNA is not methylated and serves to assess bisulfite conversion efficiency. Hence, the column “Unmethylated (percent)” indicates bisulfite conversion efficiency.

| Parental | F1 | Context | Methylated (percent) | Unmethylated (percent) | Number of mC | Number of C | Total Cytosines | Clones analyzed |
|---|---|---|---|---|---|---|---|---|
| SG292 | SG335 | All | 1.7% | 98.3% | 13 | 732 | 745 | 25 |
| SG292 | SG335 | CG | 1.4% | 98.6% | 1 | 72 | 73 | 25 |
| SG292 | SG335 | CHG | 1.3% | 98.7% | 2 | 147 | 149 | 25 |
| SG292 | SG335 | CHH | 1.9% | 98.1% | 10 | 513 | 523 | 25 |
| SG292 | SG369 | All | 6.2% | 93.8% | 11 | 166 | 177 | 6 |
| SG292 | SG369 | CG | 6.3% | 93.8% | 1 | 15 | 16 | 6 |
| SG292 | SG369 | CHG | 5.6% | 94.4% | 2 | 34 | 36 | 6 |
| SG292 | SG369 | CHH | 6.4% | 93.6% | 8 | 117 | 125 | 6 |
| SG298 | SG350 | All | 2.6% | 97.5% | 13 | 497 | 510 | 17 |
| SG298 | SG350 | CG | 0.0% | 100.0% | 0 | 51 | 51 | 17 |
| SG298 | SG350 | CHG | 4.9% | 95.1% | 5 | 97 | 102 | 17 |
| SG298 | SG350 | CHH | 2.2% | 97.8% | 8 | 349 | 357 | 17 |
| SG298 | SG362 | All | 4.7% | 95.3% | 21 | 427 | 448 | 15 |
| SG298 | SG362 | CG | 2.3% | 97.7% | 1 | 43 | 44 | 15 |
| SG298 | SG362 | CHG | 4.5% | 95.5% | 4 | 85 | 89 | 15 |
| SG298 | SG362 | CHH | 5.1% | 94.9% | 16 | 299 | 315 | 15 |
| SG307 | SG355 | All | 0.7% | 99.3% | 3 | 417 | 420 | 14 |
| SG307 | SG355 | CG | 0.0% | 100.0% | 0 | 42 | 42 | 14 |
| SG307 | SG355 | CHG | 0.0% | 100.0% | 0 | 84 | 84 | 14 |
| SG307 | SG355 | CHH | 1.0% | 99.0% | 3 | 291 | 294 | 14 |
| SG307 | SG356 | All | 2.8% | 97.2% | 11 | 378 | 389 | 13 |
| SG307 | SG356 | CG | 2.6% | 97.4% | 1 | 38 | 39 | 13 |
| SG307 | SG356 | CHG | 2.6% | 97.4% | 2 | 76 | 78 | 13 |
| SG307 | SG356 | CHH | 2.9% | 97.1% | 8 | 264 | 272 | 13 |
| SG310 | SG358 | All | 1.1% | 98.9% | 3 | 267 | 270 | 9 |
| SG310 | SG358 | CG | 0.0% | 100.0% | 0 | 27 | 27 | 9 |
| SG310 | SG358 | CHG | 0.0% | 100.0% | 0 | 54 | 54 | 9 |
| SG310 | SG358 | CHH | 1.6% | 98.4% | 3 | 186 | 189 | 9 |
| SG314 | SG346 | All | 1.2% | 98.8% | 4 | 326 | 330 | 11 |
| SG314 | SG346 | CG | 0.0% | 100.0% | 0 | 33 | 33 | 11 |
| SG314 | SG346 | CHG | 1.5% | 98.5% | 1 | 65 | 66 | 11 |
| SG314 | SG346 | CHH | 1.3% | 98.7% | 3 | 228 | 231 | 11 |
| SG314 | SG357 | All | 4.5% | 95.5% | 20 | 427 | 447 | 15 |
| SG314 | SG357 | CG | 4.7% | 95.3% | 2 | 41 | 43 | 15 |
| SG314 | SG357 | CHG | 4.4% | 95.6% | 4 | 86 | 90 | 15 |
| SG314 | SG357 | CHH | 4.5% | 95.5% | 14 | 300 | 314 | 15 |
| SG330 | SG349 | All | 3.4% | 96.7% | 14 | 404 | 418 | 14 |
| SG330 | SG349 | CG | 4.9% | 95.1% | 2 | 39 | 41 | 14 |
| SG330 | SG349 | CHG | 1.2% | 98.8% | 1 | 82 | 83 | 14 |
| SG330 | SG349 | CHH | 3.7% | 96.3% | 11 | 283 | 294 | 14 |
| SG330 | SG353 | All | 2.1% | 97.9% | 5 | 235 | 240 | 8 |
| SG330 | SG353 | CG | 4.2% | 95.8% | 1 | 23 | 24 | 8 |
| SG330 | SG353 | CHG | 4.2% | 95.8% | 2 | 46 | 48 | 8 |
| SG330 | SG353 | CHH | 1.2% | 98.8% | 2 | 166 | 168 | 8 |
| SG330 | SG359 | All | 2.4% | 97.6% | 13 | 520 | 533 | 18 |
| SG330 | SG359 | CG | 1.9% | 98.1% | 1 | 51 | 52 | 18 |
| SG330 | SG359 | CHG | 0.0% | 100.0% | 0 | 107 | 107 | 18 |
| SG330 | SG359 | CHH | 3.2% | 96.8% | 12 | 362 | 374 | 18 |

**Table S4.** Methylation analysis by Sanger sequencing after bisulfite sequencing. Target: nopaline synthase promoter (*nosP*).

| Parental | F1 | Context | Methylated (percent) | Unmethylated (percent) | Number of mC | Number of C | Total Cytosines | Clones analyzed |
|---|---|---|---|---|---|---|---|---|
| SG292 | SG335 | All | 7.7% | 92.3% | 93 | 1111 | 1204 | 17 |
| SG292 | SG335 | CG | 14.7% | 85.3% | 50 | 289 | 339 | 17 |
| SG292 | SG335 | CHG | 8.4% | 91.6% | 20 | 218 | 238 | 17 |
| SG292 | SG335 | CHH | 3.7% | 96.3% | 23 | 604 | 627 | 17 |
| SG292 | SG337 | All | 5.9% | 94.1% | 67 | 1064 | 1131 | 16 |
| SG292 | SG337 | CG | 14.4% | 85.6% | 46 | 274 | 320 | 16 |
| SG292 | SG337 | CHG | 4.0% | 96.0% | 9 | 214 | 223 | 16 |
| SG292 | SG337 | CHH | 2.0% | 98.0% | 12 | 576 | 588 | 16 |
| SG292 | SG369 | All | 11.2% | 88.8% | 71 | 562 | 633 | 9 |
| SG292 | SG369 | CG | 21.1% | 78.9% | 38 | 142 | 180 | 9 |
| SG292 | SG369 | CHG | 9.6% | 90.4% | 12 | 113 | 125 | 9 |
| SG292 | SG369 | CHH | 6.4% | 93.6% | 21 | 307 | 328 | 9 |
| SG298 | SG350 | All | 10.2% | 89.9% | 172 | 1522 | 1694 | 24 |
| SG298 | SG350 | CG | 23.9% | 76.1% | 114 | 363 | 477 | 24 |
| SG298 | SG350 | CHG | 7.8% | 92.2% | 26 | 307 | 333 | 24 |
| SG298 | SG350 | CHH | 3.6% | 96.4% | 32 | 852 | 884 | 24 |
| SG298 | SG362 | All | 10.8% | 89.2% | 107 | 884 | 991 | 15 |
| SG298 | SG362 | CG | 24.2% | 75.8% | 69 | 216 | 285 | 15 |
| SG298 | SG362 | CHG | 5.6% | 94.4% | 11 | 184 | 195 | 15 |
| SG298 | SG362 | CHH | 5.3% | 94.7% | 27 | 484 | 511 | 15 |
| SG307 | SG342 | All | 8.9% | 91.1% | 77 | 785 | 862 | 13 |
| SG307 | SG342 | CG | 18.6% | 81.4% | 46 | 201 | 247 | 13 |
| SG307 | SG342 | CHG | 2.4% | 97.6% | 4 | 164 | 168 | 13 |
| SG307 | SG342 | CHH | 6.0% | 94.0% | 27 | 420 | 447 | 13 |
| SG307 | SG355 | All | 4.4% | 95.6% | 84 | 1811 | 1895 | 27 |
| SG307 | SG355 | CG | 8.6% | 91.4% | 46 | 491 | 537 | 27 |
| SG307 | SG355 | CHG | 3.2% | 96.8% | 12 | 361 | 373 | 27 |
| SG307 | SG355 | CHH | 2.6% | 97.4% | 26 | 959 | 985 | 27 |
| SG307 | SG356 | All | 10.7% | 89.3% | 187 | 1556 | 1743 | 28 |
| SG307 | SG356 | CG | 23.9% | 76.1% | 119 | 378 | 497 | 28 |
| SG307 | SG356 | CHG | 7.0% | 93.0% | 24 | 317 | 341 | 28 |
| SG307 | SG356 | CHH | 4.9% | 95.1% | 44 | 861 | 905 | 28 |
| SG310 | SG340 | All | 8.2% | 91.8% | 97 | 1082 | 1179 | 17 |
| SG310 | SG340 | CG | 14.5% | 85.5% | 48 | 284 | 332 | 17 |
| SG310 | SG340 | CHG | 9.9% | 90.1% | 23 | 210 | 233 | 17 |
| SG310 | SG340 | CHH | 4.2% | 95.8% | 26 | 588 | 614 | 17 |
| SG310 | SG358 | All | 3.4% | 96.6% | 41 | 1163 | 1204 | 17 |
| SG310 | SG358 | CG | 6.8% | 93.2% | 23 | 315 | 338 | 17 |
| SG310 | SG358 | CHG | 2.5% | 97.5% | 6 | 232 | 238 | 17 |
| SG310 | SG358 | CHH | 1.9% | 98.1% | 12 | 616 | 628 | 17 |
| SG314 | SG346 | All | 23.4% | 76.6% | 432 | 1414 | 1846 | 26 |
| SG314 | SG346 | CG | 52.3% | 47.7% | 272 | 248 | 520 | 26 |
| SG314 | SG346 | CHG | 24.7% | 75.3% | 90 | 274 | 364 | 26 |
| SG314 | SG346 | CHH | 7.3% | 92.7% | 70 | 892 | 962 | 26 |
| SG314 | SG354 | All | 24.8% | 75.2% | 176 | 534 | 710 | 10 |
| SG314 | SG354 | CG | 55.0% | 45.0% | 110 | 90 | 200 | 10 |
| SG314 | SG354 | CHG | 27.1% | 72.9% | 38 | 102 | 140 | 10 |
| SG314 | SG354 | CHH | 7.6% | 92.4% | 28 | 342 | 370 | 10 |
| SG314 | SG357 | All | 24.0% | 76.0% | 419 | 1329 | 1748 | 25 |
| SG314 | SG357 | CG | 56.7% | 43.3% | 281 | 215 | 496 | 25 |
| SG314 | SG357 | CHG | 22.4% | 77.6% | 77 | 267 | 344 | 25 |
| SG314 | SG357 | CHH | 6.7% | 93.3% | 61 | 847 | 908 | 25 |
| SG330 | SG349 | All | 12.2% | 87.8% | 285 | 2048 | 2333 | 36 |
| SG330 | SG349 | CG | 26.9% | 73.1% | 177 | 480 | 657 | 36 |
| SG330 | SG349 | CHG | 9.9% | 90.1% | 46 | 418 | 464 | 36 |
| SG330 | SG349 | CHH | 5.1% | 94.9% | 62 | 1150 | 1212 | 36 |
| SG330 | SG353 | All | 7.0% | 93.1% | 38 | 509 | 547 | 8 |
| SG330 | SG353 | CG | 16.6% | 83.4% | 26 | 131 | 157 | 8 |
| SG330 | SG353 | CHG | 4.6% | 95.4% | 5 | 103 | 108 | 8 |
| SG330 | SG353 | CHH | 2.5% | 97.5% | 7 | 275 | 282 | 8 |
| SG330 | SG359 | All | 13.2% | 86.8% | 280 | 1840 | 2120 | 32 |
| SG330 | SG359 | CG | 30.5% | 69.5% | 183 | 417 | 600 | 32 |
| SG330 | SG359 | CHG | 10.2% | 89.8% | 43 | 377 | 420 | 32 |
| SG330 | SG359 | CHH | 4.9% | 95.1% | 54 | 1046 | 1100 | 32 |

**Table S5.** Seedling viability of 99 transgenic lines.

| Line | Transgene insertion site (Chr1) | Genomic bin (300kb) | Viability score |
|---|---|---|---|
| SG535 | 163419 | 1 | 1 |
| SG530 | 389812 | 300001 | 1.5 |
| SG521 | 880496 | 600001 | 1 |
| SG564 | 982506 | 900001 | 1 |
| SG519 | 1431190 | 1200001 | 2 |
| SG524 | 1569192 | 1500001 | 4 |
| SG346 | 2085299 | 1800001 | 1 |
| SG539 | 2249133 | 2100001 | 4.5 |
| SG489 | 2498836 | 2400001 | 10 |
| SG341 | 2952334 | 2700001 | 6 |
| SG555 | 3064124 | 3000001 | 1.5 |
| SG511 | 3442327 | 3300001 | 2 |
| SG561 | 4084162 | 3900001 | 3 |
| SG554 | 4286204 | 4200001 | 4 |
| SG505 | 4694372 | 4500001 | 1.5 |
| SG568 | 4927011 | 4800001 | 1.5 |
| SG547 | 5138334 | 5100001 | 7 |
| SG498 | 5578431 | 5400001 | 6 |
| SG315 | 6009988 | 6000001 | 1.5 |
| SG546 | 6528984 | 6300001 | 1 |
| SG531 | 6765667 | 6600001 | 3.5 |
| SG309 | 7042672 | 6900001 | 3 |
| SG335 | 7458618 | 7200001 | 8 |
| SG556 | 7232006 | 7200001 | 1 |
| SG548 | 7527079 | 7500001 | 2 |
| SG313 | 8002649 | 7800001 | 3 |
| SG518 | 8317396 | 8100001 | 2.5 |
| SG549 | 8584039 | 8400001 | 7 |
| SG516 | 8951202 | 8700001 | 6 |
| SG558 | 9076048 | 9000001 | 3 |
| SG553 | 9481532 | 9300001 | 2 |
| SG520 | 9611103 | 9600001 | 2 |
| SG565 | 9907202 | 9900001 | 4.5 |
| SG502 | 10407379 | 10200001 | 2.5 |
| SG522 | 10790022 | 10500001 | 2 |
| SG311 | 11012274 | 10800001 | 2 |
| SG506 | 11374893 | 11100001 | 2 |
| SG513 | 11575320 | 11400001 | 2 |
| SG570 | 11993298 | 11700001 | 2 |
| SG503 | 12087580 | 12000001 | 1 |
| SG544 | 12398435 | 12300001 | 1 |
| SG494 | 12842158 | 12600001 | 1 |
| SG340 | 12964622 | 12900001 | 8 |
| SG532 | 13277274 | 13200001 | 2 |
| SG534 | 13669441 | 13500001 | 7 |
| SG304 | 14056029 | 13800001 | 1 |
| SG537 | 14313049 | 14100001 | 1 |
| SG508 | 14543277 | 14400001 | 1.5 |
| SG571 | 15179141 | 15000001 | 8 |
| SG499 | 15485350 | 15300001 | 8.5 |
| SG491 | 15849233 | 15600001 | 2 |
| SG316 | 15951543 | 15900001 | 2 |
| SG496 | 16266641 | 16200001 | 1 |
| SG509 | 16646732 | 16500001 | 1 |
| SG373 | 17053813 | 16800001 | 9 |
| SG487 | 17165127 | 17100001 | 5 |
| SG512 | 17603504 | 17400001 | 7.5 |
| SG351 | 17995869 | 17700001 | 8 |
| SG541 | 18284335 | 18000001 | 1 |
| SG540 | 18473908 | 18300001 | 1 |
| SG517 | 18876837 | 18600001 | 1 |
| SG567 | 19136251 | 18900001 | 2 |
| SG507 | 19201789 | 19200001 | 2 |
| SG306 | 20068163 | 19800001 | 1 |
| SG488 | 20286882 | 20100001 | 2 |
| SG528 | 20433912 | 20400001 | 8 |
| SG526 | 20715563 | 20700001 | 1 |
| SG308 | 21043366 | 21000001 | 6.5 |
| SG504 | 21408623 | 21300001 | 1 |
| SG533 | 21839858 | 21600001 | 6 |
| SG305 | 22043881 | 21900001 | 7 |
| SG563 | 22043829 | 21900001 | 1.5 |
| SG543 | 22271226 | 22200001 | 1 |
| SG350 | 22625361 | 22500001 | 9.5 |
| SG368 | 22642823 | 22500001 | 8 |
| SG566 | 22911205 | 22800001 | 1 |
| SG486 | 23230956 | 23100001 | 1.5 |
| SG523 | 23577333 | 23400001 | 2 |
| SG569 | 23795050 | 23700001 | 1.5 |
| SG500 | 24054143 | 24000001 | 2 |
| SG545 | 24489780 | 24300001 | 3 |
| SG527 | 24861876 | 24600001 | 5.5 |
| SG536 | 25198182 | 24900001 | 2 |
| SG515 | 25474140 | 25200001 | 1 |
| SG551 | 25746736 | 25500001 | 7 |
| SG550 | 26378794 | 26100001 | 5.5 |
| SG352 | 26951796 | 26700001 | 3.5 |
| SG497 | 27437928 | 27300001 | 8 |
| SG538 | 27752400 | 27600001 | 6.5 |
| SG495 | 28151924 | 27900001 | 1 |
| SG525 | 28327886 | 28200001 | 3 |
| SG559 | 28705527 | 28500001 | 2 |
| SG303 | 29044174 | 28800001 | 7 |
| SG560 | 28960387 | 28800001 | 1 |
| SG510 | 29148256 | 29100001 | 1.5 |
| SG514 | 29610494 | 29400001 | 3.5 |
| SG529 | 29976725 | 29700001 | 3 |
| SG318 | 30038603 | 30000001 | 1.5 |
| SG493 | 30405432 | 30300001 | 5 |

**Table S6.** Viability analysis by digital image analysis. Images acquired from 14-day-old seedlings were used to determine total area covered by seedling tissue and mean grey value of the green channel using the ImageJ image analysis software.

| F1 | Area | Mean value green channel | Area x Mean | Max | Maternal line | Paternal line | Maternal phenotype | Paternal phenotype | Progeny class |
|---|---|---|---|---|---|---|---|---|---|
| SG381 | 2715218 | 203.0 | 551267995.3 | 255 | SG350 | SG369 | active | active | aa |
| SG383 | 2317627 | 194.7 | 451177083.3 | 253 | SG358 | SG369 | active | active | aa |
| SG387 | 2516087 | 197.9 | 497888327.7 | 255 | SG369 | SG350 | active | active | aa |
| SG389 | 1875489 | 195.7 | 366971306.2 | 255 | SG358 | SG350 | active | active | aa |
| SG399 | 1876518 | 196.6 | 368902797.1 | 252 | SG369 | SG358 | active | active | aa |
| SG400 | 2147235 | 200.0 | 429358963.4 | 251 | SG350 | SG358 | active | active | aa |
| SG417 | 2443513 | 195.4 | 477430674.5 | 255 | SG369 | SG369 | active | active | aa |
| SG418 | 2652069 | 199.8 | 530000077.2 | 251 | SG350 | SG350 | active | active | aa |
| SG420 | 2590169 | 195.7 | 506831319.1 | 248 | SG358 | SG358 | active | active | aa |
| SG393 | 2452259 | 191.8 | 470409487.2 | 253 | SG369 | SG356 | active | silenced | as |
| SG394 | 2090897 | 190.9 | 399194055.2 | 251 | SG350 | SG356 | active | silenced | as |
| SG395 | 2728205 | 200.3 | 546530394.8 | 255 | SG358 | SG356 | active | silenced | as |
| SG405 | 2817386 | 194.5 | 548052011.7 | 246 | SG369 | SG346 | active | silenced | as |
| SG406 | 2247877 | 200.1 | 449692289.6 | 255 | SG350 | SG346 | active | silenced | as |
| SG408 | 2183235 | 196.9 | 429876788.3 | 241 | SG358 | SG346 | active | silenced | as |
| SG411 | 2356989 | 197.6 | 465854161.9 | 255 | SG369 | SG359 | active | silenced | as |
| SG412 | 2291818 | 197.5 | 452654681.4 | 255 | SG350 | SG359 | active | silenced | as |
| SG414 | 2719761 | 196.0 | 532997002.7 | 253 | SG358 | SG359 | active | silenced | as |
| SG374 | 2724989 | 197.9 | 539144523.6 | 255 | SG369 | SG339 | active | WT | aw |
| SG375 | 2441905 | 194.9 | 475829608.3 | 255 | SG350 | SG339 | active | WT | aw |
| SG377 | 2472109 | 197.2 | 487517199.6 | 253 | SG358 | SG339 | active | WT | aw |
| SG382 | 2066800 | 197.3 | 407856111.6 | 247 | SG356 | SG369 | silenced | active | sa |
| SG384 | 2208681 | 200.9 | 443715178.2 | 255 | SG346 | SG369 | silenced | active | sa |
| SG385 | 2539552 | 193.9 | 492436909.7 | 250 | SG359 | SG369 | silenced | active | sa |
| SG388 | 2637839 | 196.5 | 518235125.6 | 254 | SG356 | SG350 | silenced | active | sa |
| SG390 | 1873755 | 195.7 | 366647009.6 | 255 | SG346 | SG350 | silenced | active | sa |
| SG391 | 2262987 | 194.8 | 440857023.4 | 255 | SG359 | SG350 | silenced | active | sa |
| SG401 | 2030258 | 196.6 | 399171055.6 | 247 | SG356 | SG358 | silenced | active | sa |
| SG402 | 2415199 | 201.7 | 487155299.1 | 255 | SG346 | SG358 | silenced | active | sa |
| SG403 | 2382035 | 197.1 | 469396671 | 253 | SG359 | SG358 | silenced | active | sa |
| SG396 | 949500 | 189.7 | 180085018.5 | 255 | SG346 | SG356 | silenced | silenced | ss |
| SG397 | 1183023 | 187.7 | 222025024.5 | 255 | SG359 | SG356 | silenced | silenced | ss |
| SG407 | 929527 | 189.3 | 175945518.2 | 255 | SG356 | SG346 | silenced | silenced | ss |
| SG409 | 876358 | 189.5 | 166028652.2 | 255 | SG359 | SG346 | silenced | silenced | ss |
| SG413 | 1315324 | 182.7 | 240316271.4 | 255 | SG356 | SG359 | silenced | silenced | ss |
| SG415 | 1037578 | 184.2 | 191099040.9 | 255 | SG346 | SG359 | silenced | silenced | ss |
| SG419 | 1033947 | 188.8 | 195174039.4 | 254 | SG356 | SG356 | silenced | silenced | ss |
| SG421 | 739776 | 187.9 | 139011308.2 | 255 | SG346 | SG346 | silenced | silenced | ss |
| SG422 | 764893 | 188.2 | 143949038.1 | 255 | SG359 | SG359 | silenced | silenced | ss |
| SG376 | 1052762 | 186.6 | 196416964.6 | 255 | SG356 | SG339 | silenced | WT | sw |
| SG378 | 1313899 | 184.4 | 242326334.3 | 255 | SG346 | SG339 | silenced | WT | sw |
| SG379 | 899555 | 186.2 | 167477350.8 | 254 | SG359 | SG339 | silenced | WT | sw |
| SG380 | 2699148 | 196.9 | 531373169.3 | 247 | SG339 | SG369 | WT | active | wa |
| SG386 | 2469660 | 195.1 | 481914634.4 | 255 | SG339 | SG350 | WT | active | wa |
| SG398 | 2588310 | 198.8 | 514649207.2 | 255 | SG339 | SG358 | WT | active | wa |
| SG392 | 1286285 | 187.4 | 241069103.3 | 255 | SG339 | SG356 | WT | silenced | ws |
| SG404 | 656335 | 185.6 | 121838747.7 | 255 | SG339 | SG346 | WT | silenced | ws |
| SG410 | 950428 | 189.2 | 179803869.9 | 255 | SG339 | SG359 | WT | silenced | ws |
| SG416 | 1240365 | 184.7 | 229076810 | 255 | SG339 | SG339 | WT | WT | ww |

**Table S7.** Phenotypic segregation analysis of F2 seedling populations. Seeds from four F1 self-crossed siblings were pooled and plated on media containing kanamycin. Kanamycin sensitivity was assessed in 14-day-old seedlings. *: scores were taken over from pooled data of **Table S8**. r: resistant, s: sensitive, WT: wild-type

| Selfed F1 | Sensitive offspring | Resistant offspring | Total analyzed | Not germinated | Mother | Father | Ancestral maternal genotype | Ancestral paternal genotype | Ancestral phenotype (m/p) | Replicate |
|---|---|---|---|---|---|---|---|---|---|---|
| SG399 | 3 | 49 | 52 | 0 | SG369 | SG358 | SG292 | SG310 | r/r | Rep1 |
| SG417 | 0 | 52 | 52 | 0 | SG369 | SG369 | SG292 | SG292 | r/r | Rep1 |
| SG400 | 1 | 49 | 50 | 2 | SG350 | SG358 | SG298 | SG310 | r/r | Rep1 |
| SG381 | 2 | 49 | 51 | 1 | SG350 | SG369 | SG298 | SG292 | r/r | Rep1 |
| SG418 | 1 | 47 | 48 | 4 | SG350 | SG350 | SG298 | SG298 | r/r | Rep1 |
| SG383 | 1 | 49 | 50 | 2 | SG358 | SG369 | SG310 | SG292 | r/r | Rep1 |
| SG420 | 1 | 51 | 52 | 0 | SG358 | SG358 | SG310 | SG310 | r/r | Rep1 |
| SG405 | 28 | 19 | 47 | 5 | SG369 | SG346 | SG292 | SG314 | r/s | Rep1 |
| SG393 | 11 | 37 | 48 | 4 | SG369 | SG356 | SG292 | SG307 | r/s | Rep1 |
| SG411 | 24 | 28 | 52 | 0 | SG369 | SG359 | SG292 | SG330 | r/s | Rep1 |
| SG406 | 22 | 29 | 51 | 1 | SG350 | SG346 | SG298 | SG314 | r/s | Rep1 |
| SG394 | 17 | 29 | 46 | 6 | SG350 | SG356 | SG298 | SG307 | r/s | Rep1 |
| SG412* | 129 | 65 | 194 | 14 | SG350 | SG359 | SG298 | SG330 | r/s | Rep1 |
| SG408 | 8 | 40 | 48 | 4 | SG358 | SG346 | SG310 | SG314 | r/s | Rep1 |
| SG374 | 18 | 28 | 46 | 6 | SG369 | SG339 | SG292 | SG339 | r/WT | Rep1 |
| SG375 | 13 | 36 | 49 | 3 | SG350 | SG339 | SG298 | SG339 | r/WT | Rep1 |
| SG377 | 14 | 38 | 52 | 0 | SG358 | SG339 | SG310 | SG339 | r/WT | Rep1 |
| SG388 | 9 | 40 | 49 | 3 | SG356 | SG350 | SG307 | SG298 | s/r | Rep1 |
| SG401* | 35 | 112 | 147 | 7 | SG356 | SG358 | SG307 | SG310 | s/r | Rep1 |
| SG382 | 8 | 43 | 51 | 1 | SG356 | SG369 | SG307 | SG292 | s/r | Rep1 |
| SG390 | 19 | 29 | 48 | 4 | SG346 | SG350 | SG314 | SG298 | s/r | Rep1 |
| SG402 | 20 | 32 | 52 | 0 | SG346 | SG358 | SG314 | SG310 | s/r | Rep1 |
| SG384 | 13 | 37 | 50 | 2 | SG346 | SG369 | SG314 | SG292 | s/r | Rep1 |
| SG391* | 164 | 33 | 197 | 11 | SG359 | SG350 | SG330 | SG298 | s/r | Rep1 |
| SG403* | 179 | 16 | 195 | 13 | SG359 | SG358 | SG330 | SG310 | s/r | Rep1 |
| SG385 | 24 | 28 | 52 | 0 | SG359 | SG369 | SG330 | SG292 | s/r | Rep1 |
| SG407 | 36 | 11 | 47 | 5 | SG356 | SG346 | SG307 | SG314 | s/s | Rep1 |
| SG419 | 46 | 5 | 51 | 1 | SG356 | SG356 | SG307 | SG307 | s/s | Rep1 |
| SG396 | 23 | 24 | 47 | 5 | SG346 | SG356 | SG314 | SG307 | s/s | Rep1 |
| SG415 | 50 | 0 | 50 | 2 | SG346 | SG359 | SG314 | SG330 | s/s | Rep1 |
| SG421 | 51 | 0 | 51 | 1 | SG346 | SG346 | SG314 | SG314 | s/s | Rep1 |
| SG409 | 49 | 0 | 49 | 3 | SG359 | SG346 | SG330 | SG314 | s/s | Rep1 |
| SG397 | 50 | 1 | 51 | 1 | SG359 | SG356 | SG330 | SG307 | s/s | Rep1 |
| SG422 | 49 | 0 | 49 | 3 | SG359 | SG359 | SG330 | SG330 | s/s | Rep1 |
| SG416 | 52 | 0 | 52 | 0 | SG339 | SG339 | SG339 | SG339 | s/s | Rep1 |
| SG376 | 48 | 0 | 48 | 4 | SG356 | SG339 | SG307 | SG339 | s/WT | Rep1 |
| SG378 | 52 | 0 | 52 | 0 | SG346 | SG339 | SG314 | SG339 | s/WT | Rep1 |
| SG379 | 49 | 0 | 49 | 3 | SG359 | SG339 | SG330 | SG339 | s/WT | Rep1 |
| SG386 | 17 | 34 | 51 | 1 | SG339 | SG350 | SG339 | SG298 | WT/r | Rep1 |
| SG398 | 14 | 36 | 50 | 2 | SG339 | SG358 | SG339 | SG310 | WT/r | Rep1 |
| SG380 | 7 | 43 | 50 | 2 | SG339 | SG369 | SG339 | SG292 | WT/r | Rep1 |
| SG404 | 52 | 0 | 52 | 0 | SG339 | SG346 | SG339 | SG314 | WT/s | Rep1 |
| SG392 | 47 | 0 | 47 | 5 | SG339 | SG356 | SG339 | SG307 | WT/s | Rep1 |
| SG410 | 52 | 0 | 52 | 0 | SG339 | SG359 | SG339 | SG330 | WT/s | Rep1 |
| SG436 | 1 | 47 | 48 | 4 | SG369 | SG350 | SG292 | SG298 | r/r | Rep2 |
| SG448 | 3 | 47 | 50 | 2 | SG369 | SG358 | SG292 | SG310 | r/r | Rep2 |
| SG466 | 0 | 49 | 49 | 3 | SG369 | SG369 | SG292 | SG292 | r/r | Rep2 |
| SG449 | 1 | 50 | 51 | 1 | SG350 | SG358 | SG298 | SG310 | r/r | Rep2 |
| SG467 | 51 | 0 | 51 | 1 | SG350 | SG350 | SG298 | SG298 | r/r | Rep2 |
| SG469 | 1 | 47 | 48 | 4 | SG358 | SG358 | SG310 | SG310 | r/r | Rep2 |
| SG454* | 37 | 114 | 151 | 5 | SG369 | SG346 | SG292 | SG314 | r/s | Rep2 |
| SG442 | 9 | 40 | 49 | 3 | SG369 | SG356 | SG292 | SG307 | r/s | Rep2 |
| SG460 | 17 | 34 | 51 | 1 | SG369 | SG359 | SG292 | SG330 | r/s | Rep2 |
| SG455 | 19 | 30 | 49 | 3 | SG350 | SG346 | SG298 | SG314 | r/s | Rep2 |
| SG443 | 7 | 44 | 51 | 1 | SG350 | SG356 | SG298 | SG307 | r/s | Rep2 |
| SG461 | 31 | 21 | 52 | 0 | SG350 | SG359 | SG298 | SG330 | r/s | Rep2 |
| SG457 | 8 | 40 | 48 | 4 | SG358 | SG346 | SG310 | SG314 | r/s | Rep2 |
| SG444 | 12 | 37 | 49 | 3 | SG358 | SG356 | SG310 | SG307 | r/s | Rep2 |
| SG463* | 153 | 33 | 186 | 22 | SG358 | SG359 | SG310 | SG330 | r/s | Rep2 |
| SG423 | 8 | 44 | 52 | 0 | SG369 | SG339 | SG292 | SG339 | r/WT | Rep2 |
| SG424 | 6 | 44 | 50 | 2 | SG350 | SG339 | SG298 | SG339 | r/WT | Rep2 |
| SG426 | 11 | 41 | 52 | 0 | SG358 | SG339 | SG310 | SG339 | r/WT | Rep2 |
| SG437 | 9 | 38 | 47 | 5 | SG356 | SG350 | SG307 | SG298 | s/r | Rep2 |
| SG450 | 8 | 43 | 51 | 1 | SG356 | SG358 | SG307 | SG310 | s/r | Rep2 |
| SG431 | 13 | 39 | 52 | 0 | SG356 | SG369 | SG307 | SG292 | s/r | Rep2 |
| SG439 | 18 | 30 | 48 | 4 | SG346 | SG350 | SG314 | SG298 | s/r | Rep2 |
| SG451* | 81 | 64 | 145 | 11 | SG346 | SG358 | SG314 | SG310 | s/r | Rep2 |
| SG433 | 26 | 23 | 49 | 3 | SG346 | SG369 | SG314 | SG292 | s/r | Rep2 |
| SG440* | 165 | 24 | 189 | 19 | SG359 | SG350 | SG330 | SG298 | s/r | Rep2 |
| SG452 | 38 | 14 | 52 | 0 | SG359 | SG358 | SG330 | SG310 | s/r | Rep2 |
| SG434 | 15 | 35 | 50 | 2 | SG359 | SG369 | SG330 | SG292 | s/r | Rep2 |
| SG456 | 49 | 0 | 49 | 3 | SG356 | SG346 | SG307 | SG314 | s/s | Rep2 |
| SG462 | 51 | 0 | 51 | 1 | SG356 | SG359 | SG307 | SG330 | s/s | Rep2 |
| SG468 | 3 | 49 | 52 | 0 | SG356 | SG356 | SG307 | SG307 | s/s | Rep2 |
| SG445 | 49 | 0 | 49 | 3 | SG346 | SG356 | SG314 | SG307 | s/s | Rep2 |
| SG464 | 47 | 0 | 47 | 5 | SG346 | SG359 | SG314 | SG330 | s/s | Rep2 |
| SG470 | 50 | 0 | 50 | 2 | SG346 | SG346 | SG314 | SG314 | s/s | Rep2 |
| SG458 | 49 | 2 | 51 | 1 | SG359 | SG346 | SG330 | SG314 | s/s | Rep2 |
| SG446 | 52 | 0 | 52 | 0 | SG359 | SG356 | SG330 | SG307 | s/s | Rep2 |
| SG471 | 51 | 0 | 51 | 1 | SG359 | SG359 | SG330 | SG330 | s/s | Rep2 |
| SG465 | 52 | 0 | 52 | 0 | SG339 | SG339 | SG339 | SG339 | s/s | Rep2 |
| SG425 | 48 | 4 | 52 | 0 | SG356 | SG339 | SG307 | SG339 | s/WT | Rep2 |
| SG427 | 48 | 4 | 52 | 0 | SG346 | SG339 | SG314 | SG339 | s/WT | Rep2 |
| SG428 | 52 | 0 | 52 | 0 | SG359 | SG339 | SG330 | SG339 | s/WT | Rep2 |
| SG435 | 16 | 34 | 50 | 2 | SG339 | SG350 | SG339 | SG298 | WT/r | Rep2 |
| SG447 | 12 | 40 | 52 | 0 | SG339 | SG358 | SG339 | SG310 | WT/r | Rep2 |
| SG429 | 11 | 41 | 52 | 0 | SG339 | SG369 | SG339 | SG292 | WT/r | Rep2 |
| SG453 | 50 | 0 | 50 | 2 | SG339 | SG346 | SG339 | SG314 | WT/s | Rep2 |
| SG441 | 47 | 0 | 47 | 5 | SG339 | SG356 | SG339 | SG307 | WT/s | Rep2 |
| SG459 | 52 | 0 | 52 | 0 | SG339 | SG359 | SG339 | SG330 | WT/s | Rep2 |

**Table S8.** Phenotypic segregation analysis of F2 seedling populations. Seeds from a subset of individual F1 self-crossed plants were plated on media containing kanamycin. Kanamycin sensitivity was visually assessed in 14-day-old seedlings. r: resistant, s: sensitive.

| Selfed F1 individual | Sensitive offspring | Resistant Offspring | Total analyzed | Not germinated | Mother | Father | Ancestral maternal genotype | Ancestral paternal genotype | Replicate | Ancestral phenotype (m/p) |
|---|---|---|---|---|---|---|---|---|---|---|
| SG391-1 | 50 | 1 | 51 | 1 | SG359 | SG350 | SG330 | SG298 | Rep1 | s/r |
| SG391-2 | 20 | 31 | 51 | 1 | SG359 | SG350 | SG330 | SG298 | Rep1 | s/r |
| SG391-3 | 48 | 0 | 48 | 4 | SG359 | SG350 | SG330 | SG298 | Rep1 | s/r |
| SG391-4 | 46 | 1 | 47 | 5 | SG359 | SG350 | SG330 | SG298 | Rep1 | s/r |
| SG401-1 | 11 | 40 | 51 | 1 | SG356 | SG358 | SG307 | SG310 | Rep1 | s/r |
| SG401-2 | 15 | 30 | 45 | 5 | SG356 | SG358 | SG307 | SG310 | Rep1 | s/r |
| SG401-3 | 9 | 42 | 51 | 1 | SG356 | SG358 | SG307 | SG310 | Rep1 | s/r |
| SG403-1 | 49 | 2 | 51 | 1 | SG359 | SG358 | SG330 | SG310 | Rep1 | s/r |
| SG403-2 | 36 | 11 | 47 | 5 | SG359 | SG358 | SG330 | SG310 | Rep1 | s/r |
| SG403-3 | 50 | 0 | 50 | 2 | SG359 | SG358 | SG330 | SG310 | Rep1 | s/r |
| SG403-4 | 44 | 3 | 47 | 5 | SG359 | SG358 | SG330 | SG310 | Rep1 | s/r |
| SG412-1 | 21 | 25 | 46 | 6 | SG350 | SG359 | SG298 | SG330 | Rep1 | r/s |
| SG412-2 | 44 | 5 | 49 | 3 | SG350 | SG359 | SG298 | SG330 | Rep1 | r/s |
| SG412-3 | 33 | 16 | 49 | 3 | SG350 | SG359 | SG298 | SG330 | Rep1 | r/s |
| SG412-4 | 31 | 19 | 50 | 2 | SG350 | SG359 | SG298 | SG330 | Rep1 | r/s |
| SG440-1 | 41 | 1 | 42 | 10 | SG359 | SG350 | SG330 | SG298 | Rep2 | s/r |
| SG440-2 | 49 | 0 | 49 | 3 | SG359 | SG350 | SG330 | SG298 | Rep2 | s/r |
| SG440-3 | 26 | 22 | 48 | 4 | SG359 | SG350 | SG330 | SG298 | Rep2 | s/r |
| SG440-4 | 49 | 1 | 50 | 2 | SG359 | SG350 | SG330 | SG298 | Rep2 | s/r |
| SG451-1 | 23 | 25 | 48 | 4 | SG346 | SG358 | SG314 | SG310 | Rep2 | s/r |
| SG451-2 | 39 | 10 | 49 | 3 | SG346 | SG358 | SG314 | SG310 | Rep2 | s/r |
| SG451-4 | 19 | 29 | 48 | 4 | SG346 | SG358 | SG314 | SG310 | Rep2 | s/r |
| SG452-1 | 40 | 9 | 49 | 3 | SG359 | SG358 | SG330 | SG310 | Rep2 | s/r |
| SG452-2 | 35 | 14 | 49 | 3 | SG359 | SG358 | SG330 | SG310 | Rep2 | s/r |
| SG452-3 | 41 | 7 | 48 | 4 | SG359 | SG358 | SG330 | SG310 | Rep2 | s/r |
| SG452-4 | 26 | 24 | 50 | 2 | SG359 | SG358 | SG330 | SG310 | Rep2 | s/r |
| SG454-1 | 12 | 39 | 51 | 1 | SG369 | SG346 | SG292 | SG314 | Rep2 | r/s |
| SG454-2 | 9 | 39 | 48 | 4 | SG369 | SG346 | SG292 | SG314 | Rep2 | r/s |
| SG454-3 | 16 | 36 | 52 | 0 | SG369 | SG346 | SG292 | SG314 | Rep2 | r/s |
| SG461-1 | 41 | 8 | 49 | 3 | SG350 | SG359 | SG298 | SG330 | Rep2 | r/s |
| SG461-2 | 50 | 0 | 50 | 2 | SG350 | SG359 | SG298 | SG330 | Rep2 | r/s |
| SG461-3 | 48 | 1 | 49 | 3 | SG350 | SG359 | SG298 | SG330 | Rep2 | r/s |
| SG463-1 | 38 | 13 | 51 | 1 | SG358 | SG359 | SG310 | SG330 | Rep2 | r/s |
| SG463-2 | 41 | 8 | 49 | 3 | SG358 | SG359 | SG310 | SG330 | Rep2 | r/s |
| SG463-3 | 34 | 10 | 44 | 8 | SG358 | SG359 | SG310 | SG330 | Rep2 | r/s |
| SG463-4 | 40 | 2 | 42 | 10 | SG358 | SG359 | SG310 | SG330 | Rep2 | r/s |

**Table S9.** PCR-based Genotyping of selected F2 seedling populations. PCR was performed using primers either spanning the insertion site (wild-type) or using a combination of a T-DNA specific primer and an endogenous primer (see also **Table S12**). Statistical analysis was performed using Chi-Square tests, followed by adjusting the p-values for multiple testing (Benjamini-Hochberg).

| T-DNA | homozygous | heterozygous | wild type | F1 parent | ChiSq *P* | FDR |
|---|---|---|---|---|---|---|
| SG310 | 11 | 17 | 11 | SG463_1 | 0.73 | 0.97 |
| SG310 | 10 | 19 | 11 | SG463_2 | 0.93 | 0.97 |
| SG310 | 12 | 29 | 10 | SG463_3 | 0.57 | 0.96 |
| SG310 | 20 | 18 | 12 | SG463_4 | 0.04 | 0.37 |
| SG310 | 11 | 26 | 14 | SG452_1 | 0.83 | 0.97 |
| SG310 | 11 | 23 | 16 | SG452_2 | 0.52 | 0.92 |
| SG310 | 13 | 28 | 11 | SG452_3 | 0.79 | 0.97 |
| SG310 | 10 | 31 | 10 | SG452_4 | 0.31 | 0.80 |
| SG298 | 9 | 34 | 8 | SG440_1 | 0.06 | 0.37 |
| SG298 | 18 | 25 | 9 | SG440_2 | 0.20 | 0.65 |
| SG298 | 10 | 33 | 9 | SG440_3 | 0.15 | 0.61 |
| SG298 | 11 | 32 | 8 | SG440_4 | 0.16 | 0.61 |
| SG298 | 14 | 23 | 14 | SG412_1 | 0.78 | 0.97 |
| SG298 | 13 | 25 | 14 | SG412_2 | 0.94 | 0.97 |
| SG298 | 10 | 30 | 10 | SG412_3 | 0.37 | 0.80 |
| SG298 | 14 | 31 | 6 | SG412_4 | 0.09 | 0.46 |
| SG330 | 10 | 20 | 6 | SG463_1 | 0.51 | 0.92 |
| SG330 | 7 | 15 | 8 | SG463_2 | 0.97 | 0.97 |
| SG330 | 10 | 16 | 10 | SG463_3 | 0.80 | 0.97 |
| SG330 | 13 | 22 | 9 | SG463_4 | 0.70 | 0.97 |
| SG330 | 7 | 21 | 12 | SG452_1 | 0.51 | 0.92 |
| SG330 | 8 | 20 | 4 | SG452_2 | 0.22 | 0.65 |
| SG330 | 16 | 20 | 5 | SG452_3 | 0.05 | 0.37 |
| SG330 | 4 | 32 | 12 | SG452_4 | 0.02 | 0.37 |
| SG330 | 10 | 26 | 13 | SG440_1 | 0.76 | 0.97 |
| SG330 | 14 | 30 | 7 | SG440_2 | 0.17 | 0.61 |
| SG330 | 11 | 31 | 10 | SG440_3 | 0.38 | 0.80 |
| SG330 | 12 | 28 | 12 | SG440_4 | 0.86 | 0.97 |
| SG330 | 12 | 22 | 13 | SG412_1 | 0.89 | 0.97 |
| SG330 | 20 | 18 | 12 | SG412_2 | 0.04 | 0.37 |
| SG330 | 12 | 27 | 12 | SG412_3 | 0.92 | 0.97 |
| SG330 | 9 | 30 | 11 | SG412_4 | 0.34 | 0.80 |

**Table S10.** Enrichment of *VANDAL6* and *ATLANTYS3* transposable elements in canonical *KEEs* (10 *KEEs*) and ectopic *KEEs* (10 ectopic *KEEs* in *ddm1* mutants (Feng et al., 2014)). Monte-Carlo based statistical testing revealed significant enrichment ($P < 0.0001$). Ectopic and canonical genomic *KEE* regions were defined as a 40 kb genomic region centered around the *KNOT* interaction maxima.

| TE family | in old *KEEs* | in ectopic *KEEs* | Total in genome | % within all *KEEs* |
|---|---|---|---|---|
| *VANDAL6* | 19 | 19 | 89 | 43 |
| *ATLANTYS3* | 18 | 12 | 142 | 21 |

**Table S11.** 4C amplification primers.

| Viewpoint | 4C primer 1 | 4C primer 2 | 1° RS | 2° RS | #cycles | annealing |
|---|---|---|---|---|---|---|
| SG261 | TCGAAAGCAACAGACTTTGGA | CAGTCCAAACAACTCTCGGA | HindIII | DpnII | 26 | 63°C |
| SG292 | TCTCCATGTTCGAACAACGT | TGAAAGAGAGAATCCAAGCAGAG | HindIII | DpnII | 26 | 63°C |
| SG298 | TCCGGCTTTCTCGTACTTGT | TCTTTGTTCTTTGCGATCCGA | HindIII | Taqαl | 26 | 64°C |
| SG307 | TCCCCTGTAAGCACAACAGA | GTCTTCTGATGTGGCTGCCA | HindIII | NlaIII | 26 | 63°C |
| SG310 | TCTTGCTGTCAGGTCAAGCT | TCGACATGCTACATGATAGAACT | HindIII | DpnII | 26 | 60°C |
| SG314 | ACGTCCCTTACCATCACACC | TGTTGTTGCTTGTAACCATTCTT | HindIII | DpnII | 26 | 62°C |
| SG330 | TCTGAAGCATCTCAATCTCTTGC | AGTGCAAATGTTAGGGAGAGTGA | HindIII | DpnII | 26 | 65°C |
| SG333 | TTTCTGCTCTTGCTTCTCTGA | GGTGTGAGACTTAACGCAACA | HindIII | Taqαl | 26 | 65°C |
| KEE6 | TCTCTGTTCTCAAAAGAGCAAAC | TGGCCGTTATTCAATTTCCCG | HindIII | DpnII | 29 | 65°C |

**Table S12.** Genotyping primers. RP and LP primers bind to genomic DNA sequence, whereas the RB primer binds to the right border of the *T-DNA* transgene.

| Parental | RP | LP | RB |
|---|---|---|---|
| SG261 | TCCAACACAGAAACTTGGTCC | ACTCTCTTGGACTTGCATTGC | ATTTTGCCGATTTCGGAAC |
| SG292 | AGATACAGTTTTGTCGGGTCG | TCCATGGAATAAGAGAAAAGAGC | ATTTTGCCGATTTCGGAAC |
| SG298 | ATGTGAGCTAGGCCTTAAGCC | TTTGAAACATCACAGCGAGTG | ATTTTGCCGATTTCGGAAC |
| SG307 | CCTTTGGTTATGCGAAATGAG | CAAGAAACAAAGCACTGCAAAC | ATTTTGCCGATTTCGGAAC |
| SG310 | TCTTGAACTCCATGTTCCAGG | TTTAACTTCTTGTCGCGAAGG | ATTTTGCCGATTTCGGAAC |
| SG314 | AAACCACATTGAGATTGCTGG | GAACTTGATGATTGCTCAGGG | ATTTTGCCGATTTCGGAAC |
| SG330 | CCTCGTCTTCGACATAACTGG | AACTTACCAATCCCATCGACC | ATTTTGCCGATTTCGGAAC |
| SG333 | CGGATCAGAAACTCTTGCTTG | GAGAGAACAAGCGGTGTTGAC | ATTTTGCCGATTTCGGAAC |

**Table S13.** Genomic positions of *KEEs. KEE* start and end positions are estimates based on the highest peak of interaction between *KEEs*.

| KEE ID | Chromosome | Start (bp) | End (bp) | Estimated Center (bp) |
|---|---|---|---|---|
| KEE01 | Chr1 | 7051324 | 7091324 | 7071324 |
| KEE02 | Chr2 | 4116555 | 4156555 | 4136555 |
| KEE03 | Chr3 | 1951581 | 1991581 | 1971581 |
| KEE04 | Chr3 | 3101455 | 3141455 | 3121455 |
| KEE05 | Chr3 | 16697396 | 16737396 | 16717396 |
| KEE06 | Chr3 | 22560488 | 22600488 | 22580488 |
| KEE07 | Chr4 | 11186537 | 11226537 | 11206537 |
| KEE08 | Chr4 | 15421465 | 15461465 | 15441465 |
| KEE09 | Chr5 | 4780379 | 4820379 | 4800379 |
| KEE10 | Chr5 | 10311725 | 10351725 | 10331725 |

**Table S14.** Droplet digital PCR primer and probe sequences.

| Target | Primer 1 | Primer 2 | MGB probe | Probe labeling |
|---|---|---|---|---|
| FIE | TAGCAAAGCGGTAAATATCACG | TGAAGTTCTAAGTGTGGTGAGCCA | TTCAAAATAAGATGGTTCCCCTTA | VIC |
| LYS | CAACGCTTCTAATTCGATTAGAGGT | GAGCGAAACCCGCATATCC | ACCATCGGCGATAAA | FAM |
| KAN | CGATGAATCCAGAAAAGCGG | GCTCCTGCCGAGAAAGTATCC | CGCCATGGGTCACG | FAM |

**Table S15.** Bisulfite sequencing, PCR primers.

| Target | Primer 1 | Primer 2 |
|---|---|---|
| nosP_bisSeq | GATTATTTGGATTGAGAGTGAATATGAG | TACCCRCCAATATATCCTRTCAAACACT |
| ChrC_bisSeq | AGAATAAATTAGAAAAGGTGGGGGGGGGG | CCTCCTTTRRATTTATRATTCACCTCAATC |

**Table S16.** Plant lines

| Parental | SALK_ID | NASC_ID | Chrom | Start | End | F1 |
|---|---|---|---|---|---|---|
| SG261 | SALK_126675 | N626675 | Chr1 | 22642823.00 | 22642938.00 | SG261/SG368/SG371 |
| SG292 | SALK_140062.52.80.x | N640062 | Chr1 | 7458618.00 | 7458855.00 | SG335/SG337SG369 |
| SG298 | SALK_112176.43.75.x | N612176 | Chr1 | 22625361.00 | 22625833.00 | SG350/SG361/SG362 |
| SG307 | SALK_131115.49.60.x | N631115 | Chr1 | 2952334.00 | 2952784.00 | SG342/SG355/SG356 |
| SG310 | SALK_061571.55.25.x | N561571 | Chr1 | 12964622.00 | 12965035.00 | SG340/SG358 |
| SG314 | SALK_056646.52.10.x | N556646 | Chr1 | 2085299.00 | 2085689.00 | SG346/SG354/SG357 |
| SG330 | SALK_030202.56.00.x | N530202 | Chr1 | 26951796.00 | 26951870.00 | SG352/SG353/SG359 |
| SG333 | SALK_058485.56.00.x | N558485 | Chr1 | 17053813.00 | 17054246.00 | SG366/SG367/SG373 |
| SG339 | NA | N60000 | WT | WT | WT | SG339 |

**Table S17.** Aligned read numbers and culture identifiers for all individual 4C experiments.

| Viewpoint | aligned reads | 4C_replicate | seedling population ID (F1) |
|---|---|---|---|
| KEE6 | 23189023 | KEE6_314 | SG354 |
| KEE6 | 21223451 | KEE6_330 | SG352 |
| SG261 | 9685774 | SG261_Rep1 | SG261 |
| SG261 | 1378433 | SG261_Rep3 | SG368 |
| SG261 | 6521728 | SG261_Rep5 | SG371 |
| SG261 | 5405450 | SG261_WT1 | SG339A |
| SG261 | 5784999 | SG261_WT2 | SG339B |
| SG261 | 5735150 | SG261_WT3 | SG339C |
| SG292 | 9913713 | SG292_Rep2 | SG335 |
| SG292 | 11091572 | SG292_Rep3 | SG337 |
| SG292 | 5100516 | SG292_Rep4 | SG369 |
| SG292 | 18036257 | SG292_WT1 | SG339A |
| SG292 | 17736274 | SG292_WT2 | SG339B |
| SG292 | 21479020 | SG292_WT3 | SG339C |
| SG298 | 29828444 | SG298_Rep1 | SG350 |
| SG298 | 17469105 | SG298_Rep2 | SG361 |
| SG298 | 16456891 | SG298_Rep3 | SG362 |
| SG298 | 12474547 | SG298_WT1 | SG339A |
| SG298 | 3740262 | SG298_WT2 | SG339B |
| SG298 | 4485737 | SG298_WT3 | SG339C |
| SG307 | 19553437 | SG307_Rep4 | SG342 |
| SG307 | 14139550 | SG307_Rep5 | SG355 |
| SG307 | 17357628 | SG307_Rep6 | SG356 |
| SG307 | 15779267 | SG307_WT4 | SG339A |
| SG307 | 20777186 | SG307_WT5 | SG339B |
| SG307 | 8076430 | SG307_WT6 | SG339C |
| SG310 | 5530946 | SG310_Rep1 | SG340* |
| SG310 | 5932068 | SG310_Rep3 | SG358 |
| SG310 | 10531730 | SG310_Rep6 | SG340* |
| SG310 | 1556512 | SG310_WT1 | SG339A |
| SG310 | 3414811 | SG310_WT2 | SG339B |
| SG310 | 7693095 | SG310_WT3 | SG339C |
| SG314 | 16360973 | SG314_Rep1 | SG346 |
| SG314 | 24693231 | SG314_Rep2 | SG354 |
| SG314 | 15137004 | SG314_Rep3 | SG357 |
| SG314 | 19053104 | SG314_WT1 | SG339A |
| SG314 | 7230100 | SG314_WT2 | SG339B |
| SG314 | 20987454 | SG314_WT3 | SG339C |
| SG330 | 19703711 | SG330_Rep2 | SG352 |
| SG330 | 22296675 | SG330_Rep3 | SG353 |
| SG330 | 23976923 | SG330_Rep4 | SG359 |
| SG330 | 6295082 | SG330_WT1 | SG339A |
| SG330 | 7935708 | SG330_WT2 | SG339B |
| SG330 | 26324223 | SG330_WT3 | SG339C |
| SG333 | 22901701 | SG333_Rep2 | SG366 |
| SG333 | 16657521 | SG333_Rep3 | SG367 |
| SG333 | 24261975 | SG333_Rep5 | SG373 |
| SG333 | 8652969 | SG333_WT1 | SG339A |
| SG333 | 16823014 | SG333_WT2 | SG339B |
| SG333 | 26660207 | SG333_WT3 | SG339C |

**Table S18.** mRNA sequencing alignment scores

| Seedling population ID (F1) | Parental | Aligned reads |
|---|---|---|
| SG335 | SG292 | 44403631 |
| SG337 | SG292 | 30517913 |
| SG339B | SG339 | 26522720 |
| SG339C | SG339 | 29785015 |
| SG339A | SG339 | 31035192 |
| SG340A | SG310 | 30258520 |
| SG340B | SG310 | 32174764 |
| SG342 | SG307 | 32013397 |
| SG346 | SG314 | 26445754 |
| SG350 | SG298 | 34331765 |
| SG352 | SG330 | 35278271 |
| SG353 | SG330 | 36861323 |
| SG354 | SG314 | 32518373 |
| SG355 | SG307 | 35669341 |
| SG356 | SG307 | 22944868 |
| SG357 | SG314 | 23740897 |
| SG358 | SG310 | 23620741 |
| SG359 | SG330 | 23915939 |
| SG362 | SG298 | 33649195 |
| SG366 | SG333 | 27055650 |
| SG367 | SG333 | 35093779 |
| SG369 | SG292 | 31740768 |
| SG373 | SG333 | 35911536 |

**Table S19.** sRNA sequencing alignment scores

| Seedling population ID (F1) | reads before filtering | reads after filtering | aligned reads |
|---|---|---|---|
| SG261 | 7396436 | 1405063 | 1311965 |
| SG335 | 8177634 | 1344447 | 1238617 |
| SG337 | 8334956 | 1733957 | 1613459 |
| SG339A | 5436747 | 1070151 | 1000903 |
| SG339B | 8664199 | 1577092 | 1469234 |
| SG339C | 4745576 | 1714664 | 1642939 |
| SG340 | 8472017 | 2001790 | 1876065 |
| SG342 | 8945031 | 1698366 | 1575524 |
| SG346 | 9513091 | 1945791 | 1816128 |
| SG350 | 9450975 | 1717736 | 1592063 |
| SG352 | 9623763 | 2063716 | 1944699 |
| SG353 | 5413341 | 1720001 | 1638880 |
| SG354 | 7867937 | 1583755 | 1475920 |
| SG355 | 7854309 | 1731756 | 1620929 |
| SG356 | 9235330 | 1894159 | 1765857 |
| SG357 | 7062132 | 1779930 | 1680265 |
| SG358 | 8040484 | 1527851 | 1428155 |
| SG359 | 6522046 | 994509 | 923212 |
| SG361 | 8571275 | 1559719 | 1453043 |
| SG362 | 6945679 | 1321023 | 1226693 |
| SG368 | 7695337 | 1605855 | 1503031 |
| SG369 | 10390466 | 2234459 | 2014860 |
| SG371 | 6860356 | 1399405 | 1308957 |

### Supplementary Figures

#### Figure S1

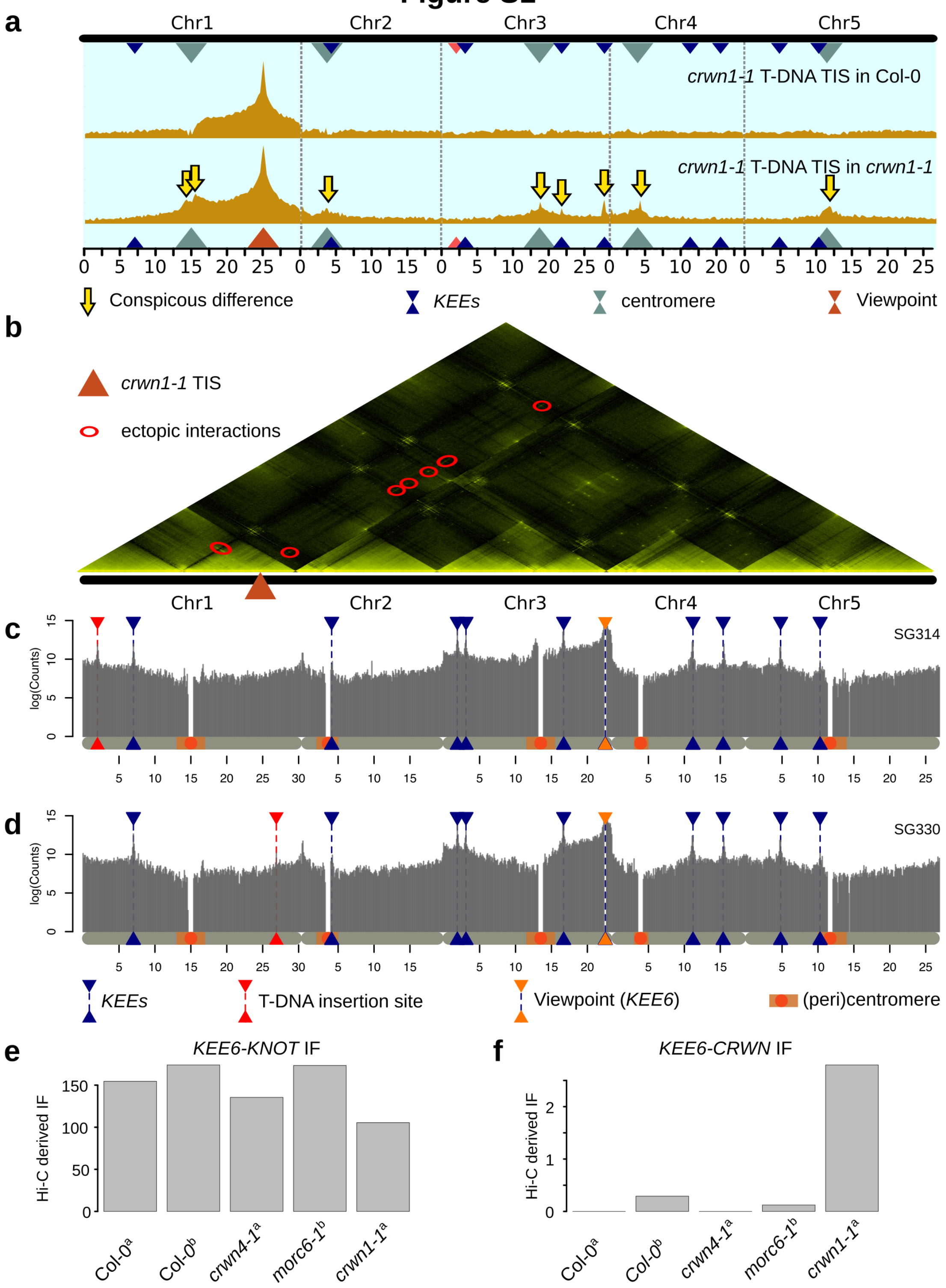


**Supplemental Figure S1. a)** Virtual 4C profile setting the *crwn1-1* TIS as a viewpoint. Top: *in silico* 4C profile extracted from the Col-0 wild-type Hi-C data set. Bottom: *in silico* 4C profile extracted from *crwn1-1* Hi-C data set (Grob et al., 2014). Bin size: 50 kb. Conspicuous differences between transgenic and wild-type virtual 4C profiles are marked with yellow arrows. **b)** Ectopic *KEEs* in *crwn1-1* shown in a Hi-C triangular plot (bin size: 100kb). **c)** 4C profile of viewpoint *KEE6* in SG314. **d)** 4C profile of viewpoint *KEE6* in SG330 **E)** Hi-C derived IFs between *KEE6* and all other *KEEs* in various genotypes. **f)** Hi-C derived IFs between *KEE6* and the *CRWN1* locus. Genotypes in **e)** and **f)** (Grob et al., 2014; Moissiard et al., 2012), bin size: 50 kb.

#### Figure S2

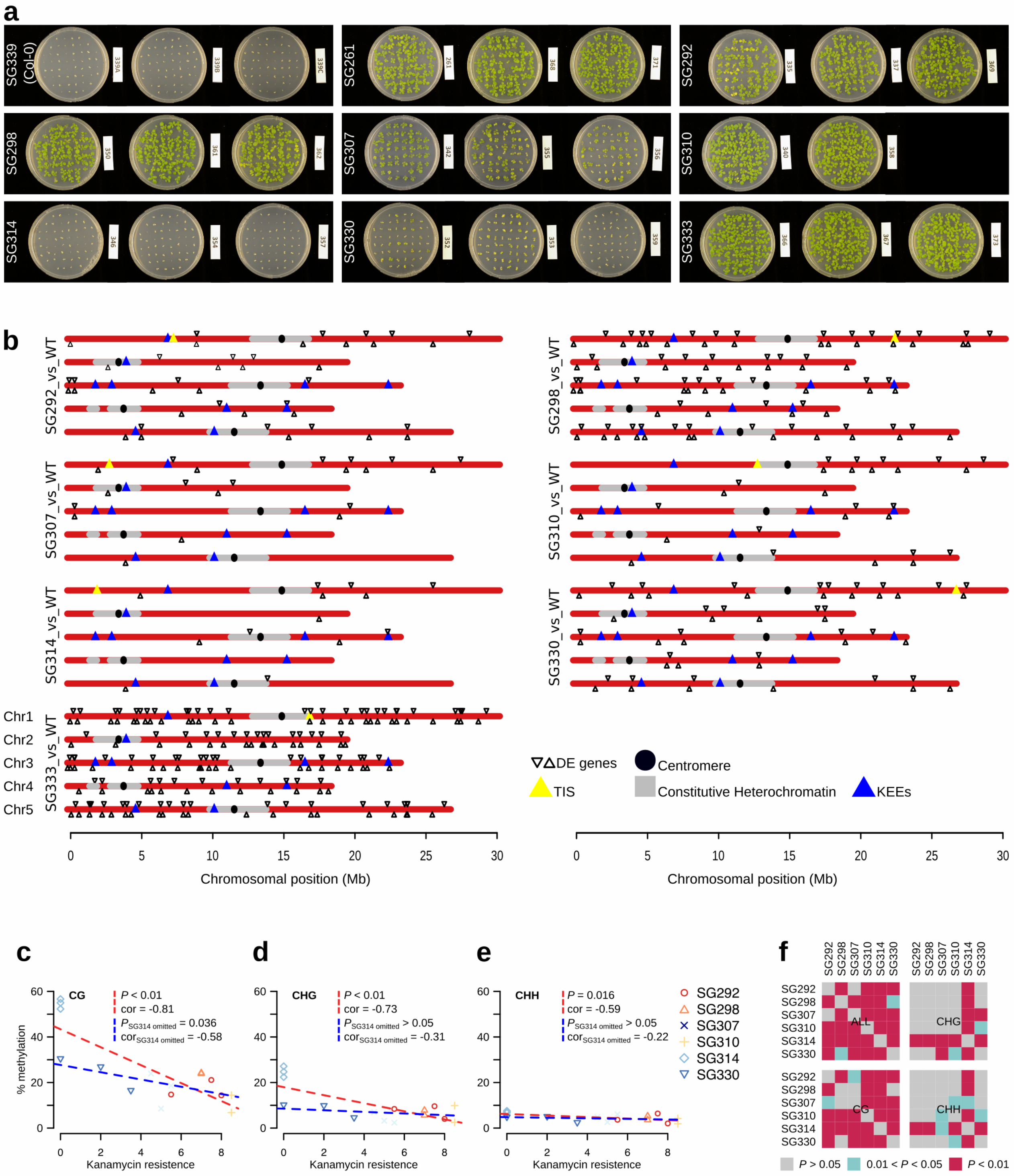


**Supplemental Figure S2.** **a)** Kanamycin resistance assay. 14-day-old seedlings grown on medium containing kanamycin. Parental line identified is indicated on the left. **b)** RNA sequencing results. Differential expression analysis between transgenic (in triplicate) and Col-0 wild-type lines (in triplicate). Differentially expressed genes (FDR < 0.05) are marked with empty black triangles. **c) - f)** Pearson's correlation analysis between kanamycin resistance phenotype and *nosP* methylation levels. **c)** CG context, **d)** CHG context, **e)** CHH context. Red dashed lines show correlation including all six transgenic lines, blue dashed lines omit SG314 in analysis. **f)** Summary of significance of cross-wise Chi-square testing between transgenic lines, split for individual mC contexts.

### Figure S3

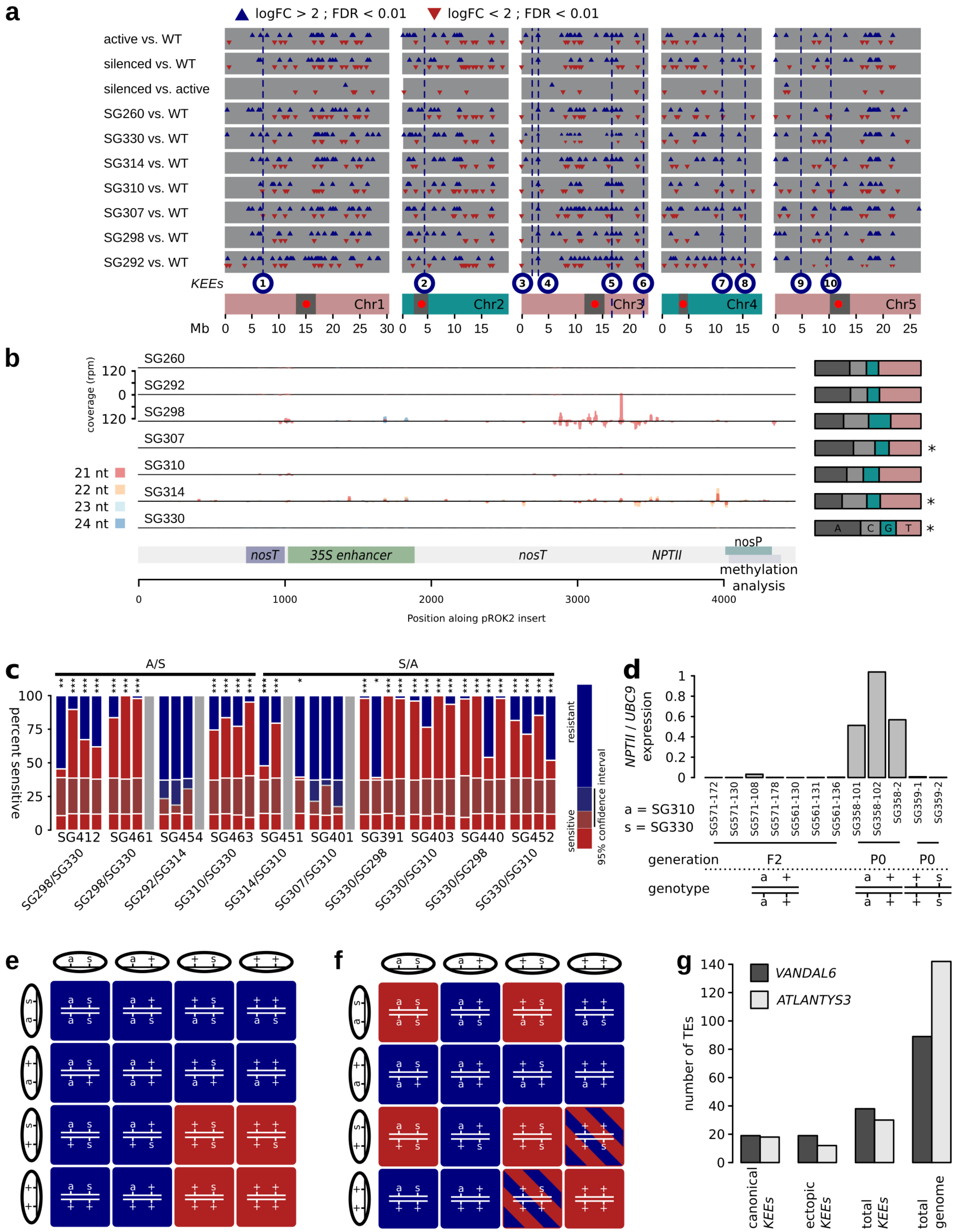


**Supplemental Figure S3. a)** sRNA-seq overview on various contrasts for which differential analysis was performed. Genomic bins (1 kb) associated with significant changes (logFC > 2; FDR < 0.01) are shown as triangles (blue, upregulation; red, downregulation). **b)** Distribution of sRNA-seq sequencing reads in 10 bp bins (21 nt - 24 nt) mapping to the vector *pROK2*. Read numbers were normalized for *pROK2* copy number. The reads of the three biological replicates were pooled. Right: Summary of first nucleotides of the reads. Asterisks mark significant deviations from equal distribution of nucleotides (Chi-Sq test, $P < 0.05$). **c)** Representation of the segregation analysis of F2 seedling populations from individual F1 mothers. Chi-square tests were performed to test for deviation from Mendelian segregation (Null-hypothesis: 0.25/0.75 (sensitive/resistant), *: $0.05 > P \geq 0.01$, **: $0.01 > P \geq 0.001$, ***: $P < 0.001$). Confidence interval indicates the range, in which Mendelian segregation cannot be rejected. A: active ancestral phenotype, S: silenced ancestral phenotype. **d)** Kanamycin expression in F2 and parental P0 plants assessed by ddPCR. **e)** Expected genotype-phenotype relationship assuming Mendelian segregation of phenotypes in the F2 generation. **f)** Expected genotype-phenotype relationship assuming a dosage effect of an interfering small RNA produced by the s locus. sRNA dosage may explain, why in the F1 generation (silenced allele in hemizygous state) no *trans*-silencing effect can be observed but only in the F2 generation. However, ratios higher than the expected ratio of kanamycin sensitive to kanamycin resistant (5/16 – 7/16) can be observed in the F2 generation (see **c)**) a: active allele, s: silenced allele, circles: maternal (top) and paternal (left) gametes, red: kanamycin sensitive phenotype, blue: kanamycin resistant phenotype. **g)** Abundance of *VANDAL6* and *ATLANTYS3* TEs in canonical and ectopic (induced by *ddm1* - (Feng et al., 2014)) *KEEs*.

### Figure S4

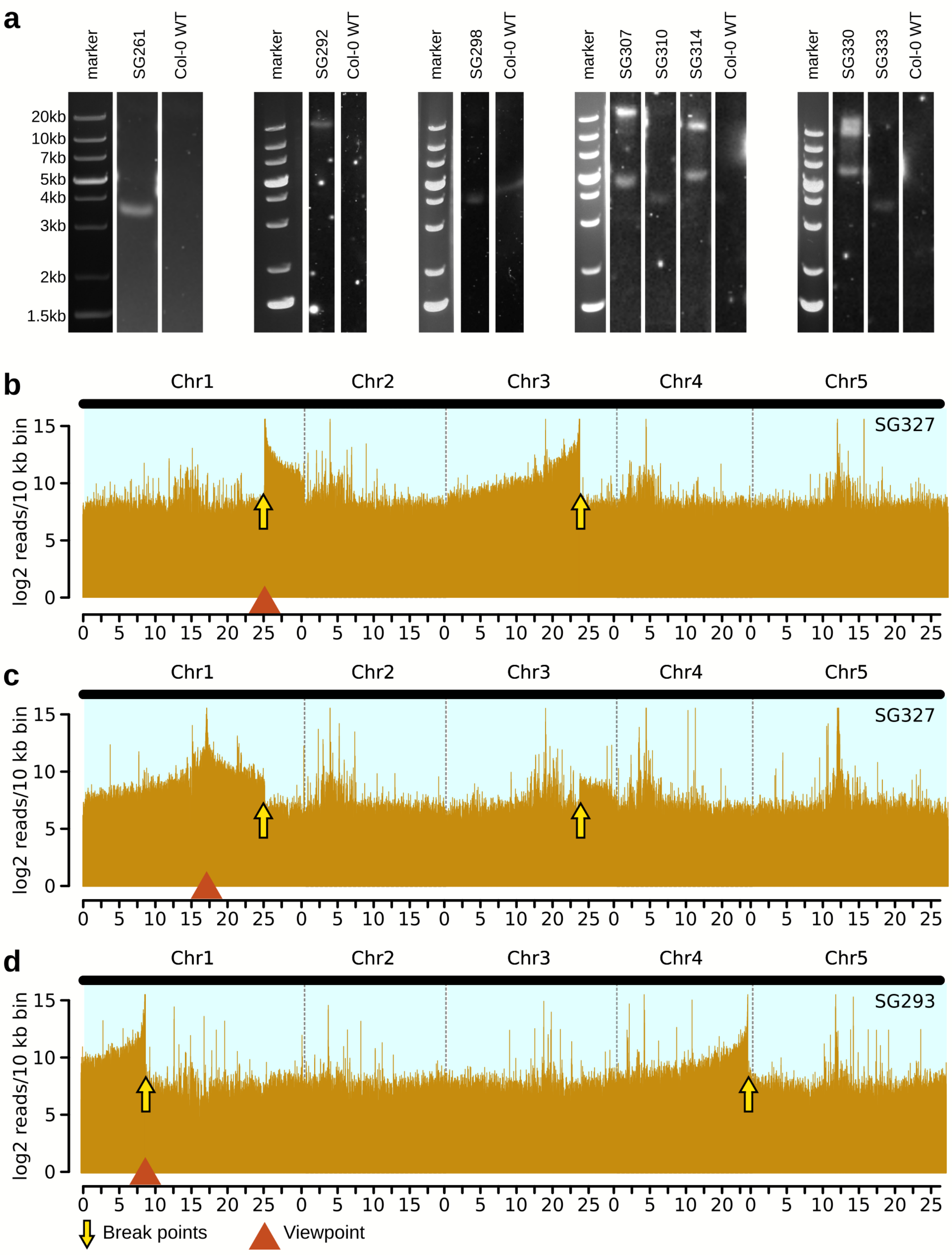


**Supplemental Figure S4. a)** Southern blot results. Single lanes have been cropped from images containing multiple lanes. DNA of plant samples non-relevant for the present study where loaded on these lanes. **b)-c)** Detection of translocations by 4C. **b)** 4C interaction frequencies from a viewpoint in close proximity of the TIS of SG327. A translocation occurred between chromosome 1 and chromosome 3. **c)** 4C interaction profile obtained from the same transgenic line as in B) with a different viewpoint. **d)** 4C interaction frequencies from a viewpoint in proximity of the TIS of SG293. A translocation occurred between chromosome 1 and chromosome 4.

#### Data Availability

4C and RNA, and sRNA sequencing data are publicly available at the Short Read Archive (SRA; https://www.ncbi.nlm.nih.gov/sra/) under accession SRP126992. Codes for data processing and analysis are available upon request.

#### Material and Methods

##### Plant Material

Seeds of transgenic *Arabidodpsis thaliana* lines were acquired through the European Arabidopsis Stock Center (NASC) (http://arabidopsis.info/) (**Table S16**). All parental lines were genotyped and homozygous individual were selected and selfed to produce F1 seeds. Plants were grown under long-day conditions (16h light, 8 hours dark, 22°C day, 18°C night). Seeds were sterilized using hypochloric acid and stratified for 2 days at 4°C. All plant material used in this study stems from 14-day-old seedlings cultivated as previously described (Grob et al., 2013). All analyzed Arabidopsis lines are in the Columbia-0 (Col-0) accession.

##### 4C experiments

Generation of 4C templates was performed as previously described (Grob et al., 2013). To minimize technical biases, we placed the 4C viewpoints adjacent to the *TIS* on endogenous DNA sequence; thus, the 4C template enrichment could be performed using identical primer pairs in transgenic and wild-type 4C samples. All 4C experiments were performed in biological triplicates. Primer sequences and restriction enzymes used in the 4C experiments are indicated in **Table S11**. 4C sequencing reads were aligned using bowtie (Langmead et al., 2009) with the parameters -a -v 0 -m 25 (no mismatches allowed, multiple alignments allowed). Alignment scores, plant lineage, and 4C replicate information are summarized in **Table S17**. Reads with multiple alignments were weighted using Rcount-multireads (Schmid and Grossniklaus, 2015). Weighted reads were mapped to individual restriction fragments using HiCdat (Schmid et al., 2015), yielding tables describing the number of reads, and thus interaction frequencies (IFs) per individual *HindIII* restriction fragment. The IFs were subsequently allocated to non-overlapping 100 kb genomic bins. IFs were further processed using edgeR (Robinson et al., 2010) to determine count data (log count per million) and to perform differential analysis between transgenic and wild-type 4C interaction profiles, comprising of 3 biological replicates each. For this, count data was normalized for library sizes (edgeR: calcNormFactors()), followed by estimating common and tag-wise dispersion (edgeR: estimateCommonDisp() and estimateTagwiseDisp()). Differential analysis was then performed using the exact test (edgeR: exactTest()). Significant differences in IFs between transgenic and WT data sets were defined by a false discovery rate (FDR) < 0.05 (using Benjamini-Hochberg adjusted *P*-values).

#### HiC data and virtual 4C analysis

Previously published Hi-C interaction data (Grob et al., 2014) were processed as previously described (Grob et al., 2014). Hi-C snapshots were taken from regions of interest using a 50 kb (**Figure 1B**) and 100 kb (**Figure 1A**) binning size. Virtual 3C and 4C analysis (see **Figure S1A** and **Figure 3F**) was performed by extracting the genomic 100 kb bin relevant to the viewpoint of interest (*crwn1-1*: Chr1 25.1 – 25.2 Mb (**Figure S1A**) or summing up Hi-C interaction frequencies between TIS and bins (100 kb) encompassing *KEEs* and pericentromeres (**Figure 3F**; *KEE1* and the pericentromere of chromosome 1 were omitted). To determine *KEE6-KNOT* contact frequencies, IFs were extracted from previously published Hi-C matrices at 50 kb resolution (Grob et al., 2014; Moissiard et al., 2012), without using distance normalization. *KEE6-KNOT* IFs were defined as contact frequencies between *KEE6* and *KEE1*, *KEE2*, *KEE3*, *KEE4*, *KEE5*, *KEE7*, *KEE8*, *KEE9*, and *KEE10*.

#### Copy number analysis

The copy numbers of inserted transgenes were assessed using Southern blot analysis and droplet digital PCR. DNA for both types of analysis was extracted from 14-day-old Arabidopsis seedlings using a MasterPure DNA purification kit (Epicentre, Madison, WIS, USA).

**a) Southern blot**

For Southern blot analysis genomic DNA was digested using the *HindIII* restriction enzyme (New England Biolabs, Ipswich, MA, USA). The digestion efficiency was analyzed on 1.5% agarose gel. Subsequently, the gel was washed for 10 min in 0.25 M HCl, followed by 15 min incubation in denaturation solution (1.5 M NaCl, 0.5 N NaOH) and 15 min incubation in neutralization solution (1.5 M NaCl, 1 M TrisHCl, pH 7.5). The fragmented and denatured DNA was then transferred to a positively charged nylon membrane (Roche, Basel, Switzerland) overnight at room temperature (RT). After rinsing the nylon membrane in 2x SSC buffer, the DNA was UV crosslinked (GS cross linker BioRad (BioRad, Hercules, CA, USA)). The membrane was then placed in a glass cylinder and incubated in 15 ml of hybridization solution (DIG Easy Hyb Granules, Roche, Basel, Switzerland) for 5 hrs at 42°C under constant rotation. Following pre-hybridization, the membrane was incubated in 15 ml of fresh hybridization solution containing 8 µl of Salmonsperm-DNA and 5 µl of digoxigenin (DIG)-labeled probe (generated by incorporation of DIG-labeled dUTP (Roche, Basel, Switzerland)) at 42°C overnight. The next day, the membrane remaining in the glass cylinder was washed 2x with W1 (2x SSC, 0.1 % SDS) for 5 min at 68°C, followed by 15 min washing in W2 (0.2x SSC, 0.1 % SDS) at 68°C, and 15 min in W3 (0.1x SSC, 0.1 % SDS). Subsequently, the membrane was transferred to a plastic tray and incubated at RT in WB (100 mM maleic acid, 150 mM NaCl, 0.3 % Tween-20, pH 7.5), followed by 30 minutes in B2 (2 g Roche Blocking Reagent (Roche, Basel, Switzerland) in 200 ml B1 (100 mM maleic acid, 150 mM NaCl, pH 7.5)). Then, 1.5 µl anti-DIG-alkaline phosphatase conjugate (Roche, Basel, Switzerland) in 50 ml B2 was added and the membrane was incubated for 30 min. The membrane was washed 3x for 40 min in WB and then incubated for 5 min in B3 (100 mM TrisHCl, 100 mM NaCl, 50 mM $MgCl_2$, pH 9.5). Finally, the membrane was overlaid with 6 ml of substrate solution (60 µl CDP Star (Roche, Basel, Switzerland) in 6 ml B3) and subsequently exposed in a trans-illuminator (Biorad Chemidoc XRS, (BioRad, Hercules, CA, USA)). Image acquisition was conducted after 10’000 sec of exposure.

**b) Droplet digital PCR**

Droplet digital PCR (ddPCR) was performed to quantify *NPTII* (and thus transgene) copy number using a Biorad QX200 Droplet Digital PCR system (BioRad, Hercules, CA, USA). The concentration of *NPTII* (transgene), *FIE* (AT3G20740; endogenous single copy gene), and *LYS* (AT5G62150; endogenous single copy gene) was assessed using 2 ng of input genomic DNA. The rounded average between *NPTII*/*FIE* and *NPTII*/*LYS* ratios was finally used to determine the *NPTII* copy number. Droplet generation was performed according to the manufacturer’s protocol. Following droplet generation, the templates were amplified in a T100 thermal cycler (BioRad, Hercules, CA, USA). Fluoresence reads of the individual droplets were analyzed using Quanta Soft v1.7 (BioRad, Hercules, CA, USA). For each sample and probe, experiments were performed in technical duplicates. Primer and probe sequence information are shown in **Table S14**. Probes were custom designed and acquired from Life Technologies (Life Technologies, ThermoFisher Scientific, Waltham, MA, USA).

#### mRNA sequencing

RNA was extracted from 14-day-old Arabidopsis seedlings using RNeasy Plant Mini Kit (Qiagen, Venlo, Netherlands). RNA was extracted from three F1 seedling populations per following parental plant lines: SG339 (wild type), SG292, SG298, SG307, SG310, SG314, SG330, and SG333. After library preparation using the Illumina Stranded mRNA RNA-seq protocol, total RNA was subjected to Illumina RNA sequencing (RNAseq). RNAseq reads were aligned using the subjunc (Liao et al., 2013) RNA sequencing reads alignment program. The numbers of valid alignments are shown in **Table S18.** Aligned RNAseq reads were then weighted and mapped to individual transcriptional units (genes, TEs) using Rcount (Schmid and Grossniklaus, 2015). The preprocessed transcription data was analyzed using the edgeR (Robinson et al., 2010) differential expression (DE) analysis program. DE was analyzed for two types of contrasts: i) individual parental transgenic lines (using three F1 seedling populations) versus Col-0 WT, and ii) all combined transgenic lines versus WT. To analyze DE, we chose a standard approach using general linearized models. After estimating common and trended (edgeR: estimateGLMCommonDisp() and estimateGLMTrendedDisp()) dispersion we applied a gene-wise negative binomial generalized linear model (edgeR: glmfit(), followed by glmLRT()) to assess DE. *P*-values were adjusted according to Benjamini-Hochberg and DE genes exhibiting adjusted *P*-values < 0.05 were scored as significant.

#### sRNA-seq analysis

Total RNA was extracted from 14-day-old *Arabidopsis* seedlings using the mirVana miRNA isolation kit (Ambion, ThermoFisher Scientific, Waltham, MA, USA). RNA was extracted from three F1 seedling populations of the following parental lines: SG339 (Col-0 wild-type; SG339A, SG339B, SG339C), SG260 (SG261, SG368, SG371), SG292 (SG335, SG337, SG369), SG298 (SG350, SG361, SG362), SG307 (SG342, SG355, SG356), SG310 (SG310, SG340), SG314 (SG346, SG355, SG356), and SG330 (SG352, SG353. SG359).
Total RNA was ligated to Illumina sequencing adapters, size selected, and subsequently sequenced on Illumina HighSeq 2500. The adapters of Illumina sequencing reads were trimmed using cutadapt (parameters: -m 17 –q 20; adapter sequence: TGGAATTCTCGGGTGCCAAGGAACTCCAGTCAC). Subsequently, the trimmed reads were filtered by aligning them against regions encompassing rRNA genes (10 kb surrounding them, Chr2:1..10000, Chr3:14194000..14204000), tRNA genes, as well as chloroplast and mitochondrial sequences. The unaligned reads were size selected (17 – 30 bp) using an awk command, and subsequently aligned to the *Arabidopsis* reference genome (TAIR10) using bowtie with the following parameters: bowtie –v 2 –best –m 10000 (allowing two mismatches and up to 10,000 equally best alignments). The aligned sequencing reads were corrected for multiple alignment using Rcount-multireads, and subsequently binned into 500 bp non-overlapping genomic bins. Differential analysis was performed using edgeR with the same parameters as described for differential RNA-seq analyses. To find genomic features associated with differential 500 bp bins, bedtools (Quinlan and Hall, 2010) intersect has been employed.

#### *NPTII* expression by ddPCR

Total RNA was extracted using standard the Trizol RNA extraction protocol, followed by RNA purification using a Direct-zol Micro Prep kit (Zymo Research, CA, USA). To remove residual DNA, RNA samples were treated with 2U of TURBO DNase following the manufacturer's protocol (Invitrogen, ThermoFisher Scientific, Waltham, MA, USA). 10 µl (ca. 1 µg) of purified RNA samples were incubated for 10 minutes at 70 °C with 1 µl oligodT, 1 µl RNase OUT (ThermoFisher Scientific, Waltham, MA, USA). Reverse transcription was performed using SuperScriptII reverse transcriptase, following the manufacturer's protocol (ThermoFisher Scientific, Waltham, MA, USA). ddPCR was performed amplifying both *NPTII* transcripts and *UBC9* transcripts as internal control to normalize for different amount of input material.

#### Methylation analysis

DNA from 14-day-old Arabidopsis seedlings was extracted using a MasterPure DNA extraction kit (Epicentre, Madison, WIS, USA). The DNA was bisulfite converted using a EpiTect Bisulfite Kits (Qiagen, Venlo, Netherlands) according to the manufacturer's protocol. Bisulfite converted DNA was amplified using a Kapa Library Amplification Kit (Kapa Biosystems, Wilmington, MA, USA). Primer sequences are indicated in **Table S15**. PCR products (see also **Figure S3B**) were subsequently purified from an agarose gel and cloned into the pJet1.2 cloning vector (CloneJET PCR cloning Kit, ThermoFisher Scientific, Waltham, MA, USA), and transformed into DH5α *E.coli* cells. Subsequently, the extracted vectors were subjected to Sanger sequencing. The resulting sequences were trimmed and preprocessed using BISMA (http://services.ibc.uni-stuttgart.de/BDPC/BISMA/) (Rohde et al., 2010). CG, CHG, and CHH methylation levels were assessed using Kismeth (http://katahdin.mssm.edu/kismeth/revpage.pl) (Gruntman et al., 2008). Statistical analysis of the methylation data was performed as previously described (Henderson et al., 2010; Jullien et al., 2012). Methylation data of all F1 samples belonging to the same parental line were pooled and subsequently Wilson's 95% confidence interval was calculated using the R package "binom". Chloroplast DNA does not exhibit cytosine methylation, thus a region of chloroplast DNA was amplified, cloned, and sequenced to assess the bisulfite conversion efficiency (see also **Table S3 and Table S4**).

#### Kanamycin sensitivity phenotype analysis

##### a) Visual Assessment

Kanamycin sensitivity in parental lines was analyzed by visual inspection. An experimenter unaware of the experimental design was asked to judge general viability of the seedlings using previously acquired images and rate the viability between 0 (dead) and 10 (perfectly viable) (double-blind assay).

##### b) Image Data Analysis

Images were acquired from 14-day-old seedlings grown on kanamycin containing medium. The images were further processed and analyzed using the ImageJ image analysis software. The color images were split in red, green, and blue channels. All subsequent steps were conducted in the green channel images. To extract the area covered by seedlings tissue, a grey threshold was set, which has been previously empirically defined and showed the best separation between seedling tissue and background (lower threshold 160, upper threshold 255). The total area and the mean grey value within the threshold were measured. The product between total area and mean grey value (area x mean; see **Table S6**) was used to perform statistical analysis. Assuming normal distribution of the data, we performed cross-wise t-tests between progeny classes (progeny classes derived from crosses with the wild type were omitted). The *p*-values were adjusted for multiple testing using the Benjamini-Hochberg (aka FDR) algorithm. Statistical testing was conducted using R.

**c) Statistical Analysis**

Correlation between viability on kanamycin (as well as *NPTII* expression) and *TIS-KNOT* IFs was performed using the R-base cor.test() function. The slope and intercept were retrieved by employing a linear model using the R-base lm() function. To assess whether the chromosomal position may affect transgene expression, the viability score of 99 transgenic lines inserted into chromosome 1 were assessed visually. Subsequently, the along the chromosome ordered viability scores were tested for randomness using a two-sided Bartels rank test.
